# Similar, but not the same: multi-omics comparison of human valve interstitial cells and osteoblast osteogenic differentiation expanded with an estimation of data-dependent and data-independent PASEF

**DOI:** 10.1101/2024.04.03.587893

**Authors:** Arseniy Lobov, Polina Kuchur, Nadezhda Boyarskaya, Daria Perepletchikova, Ivan Taraskin, Andrei Ivashkin, Daria Kostina, Irina Khvorova, Vladimir Uspensky, Egor Repkin, Evgeny Denisov, Tatiana Gerashchenko, Rashid Tikhilov, Svetlana Bozhkova, Vitaly Karelkin, Chunli Wang, Kang Xu, Anna Malashicheva

## Abstract

Osteogenic differentiation is crucial in normal bone formation and pathological calcification, such as calcific aortic valve disease (CAVD). Understanding the proteomic and transcriptomic landscapes underlying this differentiation can unveil potential therapeutic targets for CAVD. In this study, we employed the timsTOF Pro platform to explore the proteomic profiles of valve interstitial cells (VICs) and osteoblasts during osteogenic differentiation, utilizing three data acquisition/analysis techniques: Data-Dependent Acquisition (DDA-PASEF) and Data-Independent Acquisition (DIA-PASEF) with a classic library based and machine learning-based “library-free” search (DIA-ML). RNA-seq complemented comparative proteome coverage analysis to provide a comprehensive biological reference. We reveal distinct proteomic and transcriptomic profiles between VICs and osteoblasts, highlighting specific biological processes in their osteogenic differentiation pathways. Furthermore, the study identified potential therapeutic targets for CAVD, including the differential expression of proteins such as MAOA and ERK1/2 pathway in VICs. From a technical perspective, the DIA-ML offers significant advantages and seems the method of choice for routine proteomics.

## 1. Introduction

Vessels and bone tissues exhibit intricate interconnections, where vessels provide a structural framework for bone formation and play a pivotal role in governing bone development and repair.^1^ This interrelation clarifies why, in pathological conditions, vascular tissues may undergo calcification. Calcifications can manifest in diverse forms and mechanisms throughout the body, significantly elevating health risks irrespective of location. Notably, meta-analyses have indicated that calcification in any arterial wall is linked to a 3–4-fold increase in mortality risk and cardiovascular events.^2^

*Calcific aortic valve disease (CAVD)* or calcific aortic valve stenosis is one of the most dangerous forms of vascular calcification. CAVD is a gradual condition that initiates with aortic valve thickening and gradually advances to extensive calcification. Subsequently, valve calcification in advanced stages can precipitate potentially life-threatening circulatory disorders. The prevalence of CAVD has exhibited substantial growth over time. Recent statistics indicate a rise in incident cases of CAVD globally, from 130,822 in 1990 to 589,638 in 2019.^3^ During this period, CAVD witnessed the highest surge in death rates and prevalence among the three primary valvular diseases, including rheumatic heart disease, mitral regurgitation, and CAVD.^4^ Over the past three decades, many Western nations have witnessed a prevalence increase of over 10% in CAVD.^4^ While comprehensive data is lacking for numerous resource-poor countries, the global trend underscores a universal concern.^4,5^ The existing treatment for CAVD entails surgical valve replacement using either a bio-or mechanical prosthesis, necessitating replacement every 5-15 years. It is imperative to develop therapeutic strategies that can mitigate the progression of CAVD urgently.

A fundamental lack of understanding regarding the pathogenesis of CAVD stands as a critical barrier to therapeutic interventions. While the endothelium is believed to be significant in initiating CAVD^6^, the osteogenic transformation of resident valve interstitial cells (VICs) is central to disease progression and valve calcification.^5,7^

While there are many high-quality studies on the molecular mechanisms behind VICs osteogenic differentiation^8^, there are still gaps in our understanding. To develop potential therapies, it is essential to understand the differences between VICs osteogenic trans-differentiation and normal osteogenic differentiation. Although it is generally assumed that osteogenic differentiation of VICs is similar to that of osteoblasts, there is still no empirical confirmation of this assumption because of the lack of multi-omics studies that describe the molecular mechanisms of adult osteoblast differentiation and physiology.^9^ Therefore, we aimed to perform the multi-omics comparison of molecular mechanisms of VICs and osteoblast osteogenic differentiation *in vitro*.

In omics, particularly in proteomics research, the depth of biological insights is constrained by the technical capabilities of omics methodologies. The field of mass spectrometry-based omics is advancing swiftly, with novel platforms for proteomics emerging regularly. This progress necessitates the development of specialized bioinformatics methods. However, given that the primary users of mass spectrometry-based omics frequently include biologists and clinicians who may not have expertise in mass spectrometry, conducting comparative studies is crucial. Such studies should elucidate how different platforms and data acquisition strategies impact the ability to address biological questions effectively.

The Trapped Ion Mobility Spectrometry coupled with Time-of-Flight Mass Spectrometry (TIMS-TOF) technology, exemplified by the timsTOF Pro platform from Bruker, represents a cutting-edge advancement in shotgun proteomics, showcasing superior performance capabilities.^10^ This platform supports both Data-Dependent Acquisition (DDA) and Data-Independent Acquisition (DIA) strategies, offering flexibility in experimental approaches. Moreover, various bioinformatic tools are accessible for analyzing data derived from either approach. Despite numerous studies comparing these methodologies and their corresponding analysis software using standard test samples, such as HeLa cell lysate, a significant gap exists in the literature concerning their comparison in the context of addressing specific biological inquiries in biomedical studies.^11–14^

To bridge biological and technical gaps in current knowledge, we conducted a comparative study on the molecular mechanisms underlying osteogenic differentiation in human osteoblasts isolated from adult femur bones and VICs from patients with CAVD. Our methodology combined transcriptomics with shotgun proteomics. To achieve the most comprehensive proteomic coverage and assess various PASEF techniques, we analyzed the same samples using DDA-and DIA-PASEF. For the DIA-PASEF data, a library-free search strategy utilizing DIA-NN was employed alongside a traditional library-based approach, constructing a spectral library from pre-fractionated pooled samples of each biological group under investigation.

Our findings reveal differences in molecular mechanisms of osteoblasts and VICs osteogenic differentiation, which might be used to target anti-CAVD therapy development. Specifically, we underline MAOA and ERK1/2 as fruitful targets, which are upregulated in VICs osteogenic differentiation and have been shown to have the potential for CAVD treatment before. From a technical perspective, we show that for complex samples, such as human primary cell cultures, DIA-PASEF outperforms DDA to an even greater extent than previously demonstrated with test samples. While classic DIA-PASEF with library-based data analysis appears to be a golden standard, the library-free search approach with DIA-NN proves nearly as efficient as the library-based method in recognizing differentially expressed proteins. By comparing the RNA-seq data, we observed only a minor compromise in data reliability with the DIA-NN method compared to library-based analysis.

## 2. Material and methods

A schematic representation of the study design is provided in Figure 1, followed by a detailed explanation in the text below.

**Figure 1.**
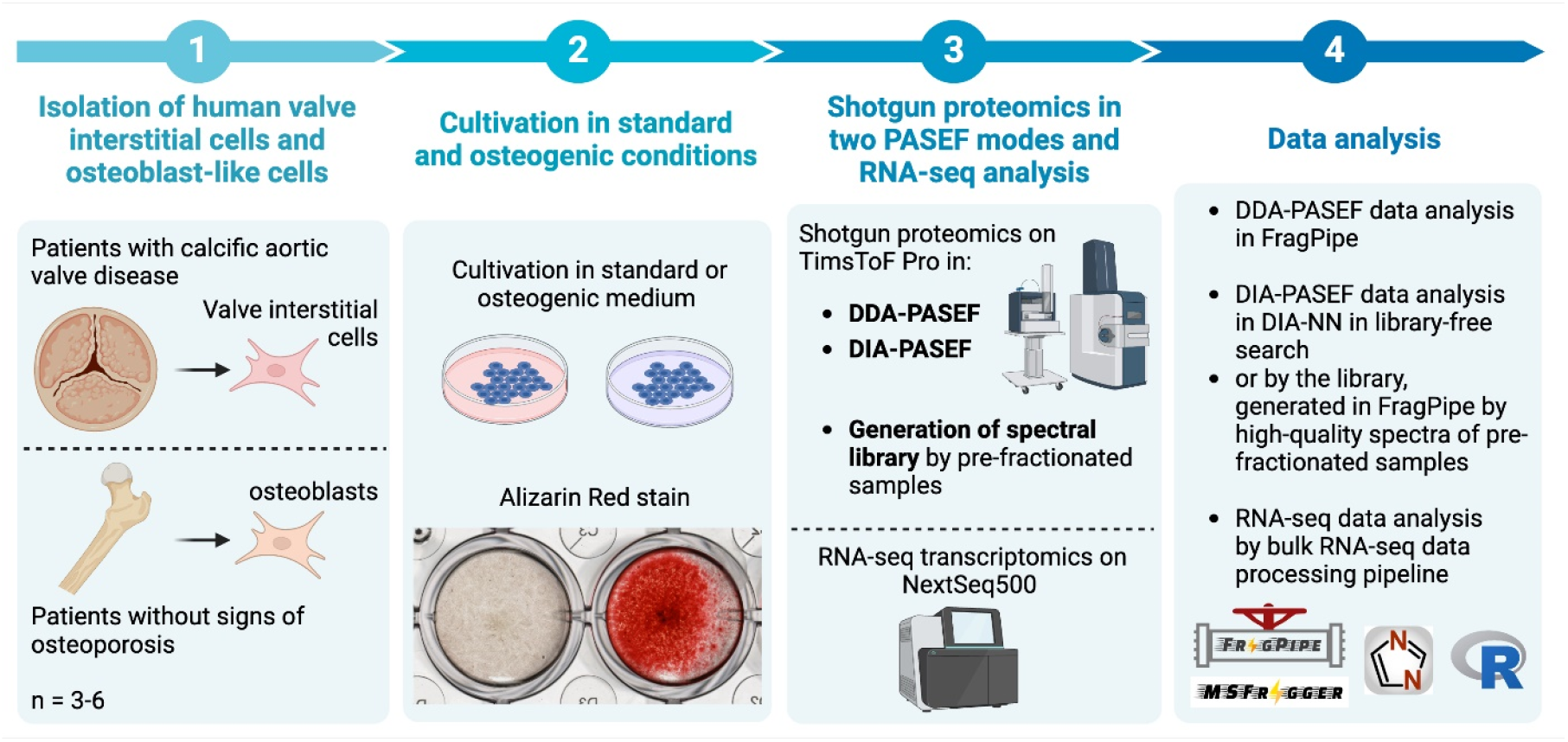
A schematic representation of the study design.

### 2.1 Cell cultures

#### Valve interstitial cell isolation

Human valve interstitial cells (VICs) were obtained from the calcified human aortic valves (n = 5), which were isolated from patients with calcific aortic valve disease (CAVD) during surgical valve replacement in the Almazov National Medical Research Centre of the Ministry of Health of the Russian Federation. The study was conducted according to the guidelines of the Declaration of Helsinki and approved by the local Ethics Committee of the Almazov Federal Medical Research Center. All patients gave informed consent. Patients with bicuspid valves were excluded.

Fragments of calcified valve were washed in PBS and treated with collagenase II (1 mg/ml; Worthington Biochemical Corporation) in PBS for 10 minutes, and then endothelial cells were scraped out. The tissues were then digested by collagenase II in a standard cultivation medium overnight at 37 °C. Tissue fragments were transferred to a cultural flask.

#### Osteoblasts isolation

Human osteoblast-like cells were obtained from the fragments of the epiphysis of bone spongy tissue of femur bone (n = 6), harvested during surgery at Vreden National Medical Research Center of Traumatology and Orthopedics. The local Ethics Committee of Vreden National Medical Research Center of Traumatology and Orthopedics approved the clinical research protocol and followed the Declaration of Helsinki principle. All patients gave informed consent. Patients with osteoporosis were excluded.

Bone fragments were washed in phosphate buffer saline (PBS) supplemented with Penicillin/Streptomycin 3 times. Then, the fragments were crushed into fragments up to 0.5 mm using carbide cutters. Cancellous bone was repeatedly washed in PBS until connective tissue and capillary were removed. Then, the fragments were incubated with 0.2% collagenase type II (Worthington Biochemical Corporation) solution in PBS for 30 min at 37°C, washed in PBS, and transferred into 0.2% collagenase type IV solution (Worthington Biochemical Corporation) in standard cultivation media and incubated for 16 hours at 37°C. Then, fragments were transferred to a culture flask, where cells were explanted for several weeks.

#### Cell culturing

Both VICs and osteoblasts were cultured in the same conditions (37 °C in 5% CO_2_) in DMEM (41966-052, Gibco) supplemented with 15% FBS (HyClone, GE Healthcare), 2 mM L-glutamine (Gibco) and penicillin/streptomycin (Gibco) until 70–80% confluence. The medium was changed twice a week. Cell cultures were routinely checked for mycoplasma contamination each week, according to Janetzko et al. 2014^15^ with minor modifications. Cells of 3-4 passages were used for the experiments.

#### Osteogenic differentiation

We used cells from the same donors for both transcriptomics and proteomics analysis. For both methods, cells were analyzed in standard cultivation and on the 10^th^ day after induction of osteogenic differentiation.

To induce osteogenic differentiation, we used a classic osteogenic medium, usually used for *in vitro* osteogenic differentiation: DMEM supplemented with 10% FBS, 2 mM L-glutamine, penicillin/streptomycin, 50 mg/ml ascorbic acid, 0.1 mM dexamethasone, and 10 mM b– glycerophosphate.^9^

Osteogenic differentiation was verified by Alizarin red stain (stain for calcium deposition) on the 21st day of osteogenic differentiation. For this purpose, in parallel with the main experiment, cells were plated in 24-well plates. After 21 days of differentiation, cells were washed with PBS, fixed in 70% ethanol for 60 min, washed twice with distilled water, and stained using Alizarin Red solution (Sigma).

### 2.2 Shotgun proteomics

#### Protein isolation

The cells were washed twice with PBS and lysed in a Petri dish with RIPA buffer (ThermoFisher Scientific) supplemented with a complete protease inhibitor cocktail (Roche). Then, the dishes were transferred to the ice for 30 min. Cell lysates were stored at −80°C before use.

The samples were sonicated for 10 min in an ultrasonic bath with ice and centrifuged (12 000 g, 20 min, 4°C). Proteins were cleaned by standard acetone precipitation (EM grade acetone; EMS). The protein pellet was resuspended in 8M Urea/50mM ammonium bicarbonate (Sigma Aldrich) and centrifuged (12 000 g, 10 min, 4°C). The protein concentration was measured by a Qubit fluorometer (ThermoFisher Scientific) using QuDye Protein Quantification Kit (Lumiprobe) according to the manufacturer’s recommendations.

#### In-solution digestion

20 μg of each sample was digested by trypsin. Sample volumes were equalized by the addition of 8M Urea/50mM ammonium bicarbonate. Disulfide bonds were reduced and alkylated by incubation for 1 h at 37 °C in 5 mM DTT (Sigma Aldrich) with subsequent incubation in 15 mM iodoacetamide for 30 min in the dark at room temperature (Sigma Aldrich). Then, the samples were diluted with seven volumes of 50 mM ammonium bicarbonate and incubated for 16 h at 37 °C with 400 ng of trypsin (ratio 1:50; “Trypsin Gold”, Promega).

After digestion, the samples were desalted by solid phase extraction in “Stage tips”, self-made from polypropylene Vertex pipette tips (200 µL; SSIbio) filled with four layers of C18 reversed-phase excised from Empore 3M C18 extraction disks according to Rappsilber et al.^16^ Desalted peptides were evaporated in Eppendorf Concentrator plus (Eppendorf) and dissolved in water/0.1% formic acid for further LC-MS/MS analysis. Each sample was analyzed in both DDA-and DIA-PASEF modes.

#### Library fractionation

To obtain a spectral library, we fractionated four pooled samples. For this purpose, we mixed all replicates across each biological group (VICs and osteoblasts in standard cultivation and after osteogenic differentiation). Then, 100 μg of each pooled sample was digested in-solution as described above and fractionated by Pierce^™^ High pH Reversed-Phase Peptide Fractionation Kit (ThermoFisher Scientific) according to manufacturer recommendations. Each fraction was desalted as described above and dissolved in water/0.1% formic acid. The peptide concentration was measured by Pierce^™^ Quantitative Fluorometric Peptide Assay (ThermoFisher Scientific) in Cary Eclipse Fluorescence Spectrometer (Agilent) according to manufacturer recommendations.

#### HPLC

Approximately 500 ng of tryptic peptides were used for shotgun proteomics analysis by nanoHPLC-MS/MS with trapped ion mobility mass spectrometry in timsTOF (Bruker). HPLC was performed in two-column separation mode with Acclaim^™^ PepMap^™^ 5 mm Trap Cartridge and Aurora Series separation column with nanoZero technology (C18, 25 cm x 75 µm ID, 1.6 µm C18) in gradient mode with 400 nL/min flow rate. Phase A was water/0.1% formic acid, phase B was acetonitrile/0.1% formic acid. The gradient was from 2% to 18% of phase B for 44 min, then to 25% of phase B for 11 min, to 37% of phase B for 5 min with subsequent wash with 95% of phase B for 17 min. The CaptiveSpray ion source was used for electrospray ionization with 1600 V of capillary voltage, 3 l/min N_2_ flow, and 180 °C source temperature. MS acquisition was performed in two different modes.

#### DDA-PASEF

MS/MS acquisition in DDA-PASEF mode was used to analyze experimental and fractionated samples for the spectral library. The analysis was performed by automatic DDA-PASEF mode with 1.1 s cycle time and 10 PASEF ramps in positive polarity with the fragmentation of ions with at least two charges in m/z range from 100 to 1700 and ion mobility range from 0.60 to 1.60 1/K0. The active exclusion was enabled with a release time of 0.4 min and reconsideration after a four-fold increase in intensity. Collision energy was 20 eV for 0.6 1/K0 and 59 eV for 1.6 1/K0. Isolation width was 2 m/z for 700 m/z and 3 m/z for 800 m/z.

#### DIA-PASEF

Analysis in DIA-PASEF mode was performed in the same m/z and ion mobility range as DDA-PASEF. The DIA analysis was performed in positive polarity with the fragmentation of ions with at least two charges in m/z range from 100 to 1700 and ion mobility range from 0.60 to 1.60 1/K0. Cycle time was 1.6 s, and it included 16 fragmentation steps with 26 m/z and 0.01-0.02 1/K0 step (tab. 1). Each step included parallel fragmentation of two fragments of the spectra with a difference in 400 m/z and 0.3 1/K0. The collision energy was 20 eV for 0.85 1/K0 and 59 eV for 1.3 1/K0.

**Table 1.** DIA-PASEF isolation list

| MS type | Cycle Id | Start IM [1/K0] | End IM [1/K0] | Start Mass [m/z] | End Mass [m/z] |
| --- | --- | --- | --- | --- | --- |
| MS1 | 0 | - | - | - | - |
| PASEF | 1 | 0.9001 | 1.2001 | 800.00 | 826.00 |
| PASEF | 1 | 0.6000 | 0.9001 | 400.00 | 426.00 |
| PASEF | 2 | 0.9201 | 1.2201 | 825.00 | 851.00 |
| PASEF | 2 | 0.6200 | 0.9201 | 425.00 | 451.00 |
| PASEF | 3 | 0.9301 | 1.2301 | 850.00 | 876.00 |
| PASEF | 3 | 0.6300 | 0.9301 | 450.00 | 476.00 |
| PASEF | 4 | 0.9500 | 1.2501 | 875.00 | 901.00 |
| PASEF | 4 | 0.6501 | 0.9500 | 475.00 | 501.00 |
| PASEF | 5 | 0.9600 | 1.2601 | 900.00 | 926.00 |
| PASEF | 5 | 0.6601 | 0.9600 | 500.00 | 526.00 |
| PASEF | 6 | 0.9800 | 1.2801 | 925.00 | 951.00 |
| PASEF | 6 | 0.6801 | 0.9800 | 525.00 | 551.00 |
| PASEF | 7 | 0.9900 | 1.2901 | 950.00 | 976.00 |
| PASEF | 7 | 0.6900 | 0.9900 | 550.00 | 576.00 |
| PASEF | 8 | 1.0101 | 1.3101 | 975.00 | 1001.00 |
| PASEF | 8 | 0.7100 | 1.0101 | 575.00 | 601.00 |
| PASEF | 9 | 1.0201 | 1.3201 | 1000.00 | 1026.01 |
| PASEF | 9 | 0.7200 | 1.0201 | 600.00 | 626.00 |
| PASEF | 10 | 1.0401 | 1.3401 | 1025.01 | 1051.01 |
| PASEF | 10 | 0.7400 | 1.0401 | 625.00 | 651.00 |
| PASEF | 11 | 1.0601 | 1.3601 | 1050.01 | 1076.01 |
| PASEF | 11 | 0.7601 | 1.0601 | 650.00 | 676.00 |
| PASEF | 12 | 1.0701 | 1.3701 | 1075.01 | 1101.01 |
| PASEF | 12 | 0.7701 | 1.0701 | 675.00 | 701.00 |
| PASEF | 13 | 1.0901 | 1.3901 | 1100.01 | 1126.01 |
| PASEF | 13 | 0.7901 | 1.0901 | 700.00 | 726.00 |
| PASEF | 14 | 1.1001 | 1.4001 | 1125.01 | 1151.01 |
| PASEF | 14 | 0.8001 | 1.1001 | 725.00 | 751.00 |
| PASEF | 15 | 1.1201 | 1.4201 | 1150.01 | 1176.01 |
| PASEF | 15 | 0.8200 | 1.1201 | 750.00 | 776.00 |
| PASEF | 16 | 1.1300 | 1.4301 | 1175.01 | 1201.01 |
| PASEF | 16 | 0.8300 | 1.1300 | 775.00 | 801.00 |

#### Data analysis

Protein identification in all cases was performed using the same *Homo sapiens* reference proteome (UP000005640, downloaded 27.06.2022) with contaminations included.

Analysis of DDA-PASEF data and spectral library generation was performed in FragPipe (v. 18.0) according to the default LFQ-MBR workflow. The search parameters were parent and fragment mass error tolerance 15 ppm, protein and peptide FDR less than 1%, protease rule – strict trypsin (cleave after KR), and two possible missed cleavage sites. Cysteine carbamidomethylation was set as a fixed modification. Methionine oxidation and acetylation of protein N-term were set as variable modifications.

DIA-PASEF data analysis was performed in DIA-NN (v. 1.8.1.) in two variants – with empirically obtained spectral library from FragPipe (DIA) or with deep-learning predicted spectral library constructed in DIA-NN (DIA-ML) with the same search parameters. The search parameters were default with minor changes: parent and fragment mass error tolerance were set in automatic interference mode and were within 13-15 ppm for all samples, protein, and peptide FDR less than 1%, protease rule – strict trypsin (cleave after KR), two possible missed cleavage sites. Cysteine carbamidomethylation was set as a fixed modification. Methionine oxidation and acetylation of protein N-term were set as variable modifications.

### 2.3 RNA-seq transcriptomics

#### RNA isolation

RNA was isolated using ExtractRNA (Eurogene) using a standard phenol/chloroform extraction procedure according to the manufacturer’s instructions. Briefly, cells were washed with PBS and lysed in a Petry dish by ExtractRNA for 5 minutes at room temperature. Then, lysates were transferred to microcentrifuge tubes and mixed with 1/5 of the volume of chloroform. The samples were incubated for 10 minutes and centrifuged (20 min, 12600g, 4°C). The aqueous phase was mixed with ½ of the initial ExtractRNA volume of ice-cold isopropanol. After incubation for 10 minutes, the samples were centrifuged (20 min, 12600g, 4°C). The precipitate was washed with 70% ethanol, air-dried at room temperature, and dissolved in water. RNA quality was assessed by agarose electrophoresis and Nanodrop spectrometry.

#### Next-generation sequencing

For RNA-seq library preparation, 500 ng of each sample was used with the “CORALL Total RNA seq Library Prep Kit” (Lexogen), which included poly-A RNA selection according to the manufacturer’s recommendation. The quality of the obtained libraries was verified by capillary electrophoresis using a 4150 TapeStation (Agilent, USA). Finally, the libraries were sequenced on the Illumina NextSeq500 platform using single-end reads. All samples were run simultaneously. Raw reads are deposited in the NCBI SRA database with BioProject identifier PRJNA947173. RNA sequencing was carried out using the equipment of the Core Facility “Medical Genomics” (Tomsk NRMC) and the Tomsk Regional Common Use Center.

#### Data analysis

Quality control of the fastq files was performed using the FASTQC and MultiQC packages. Adapter sequences were removed by the Fastp package. Then, reads were aligned to the *Homo sapiens* reference genome GRCh38 using the STAR package, with further quantification by the QoRTs package.

### 2.4. Statistical analysis

We performed statistical data analysis to compare osteoblasts and VICs at each stage (before and after osteogenic differentiation). Proteomic and transcriptomic data were analyzed separately, and then their results were compared. Statistical analysis was performed using R.

#### RNA-seq data analysis

Regarding the RNA-sequencing data analysis, we examined six samples of osteoblasts (comprising three under standard culture conditions and three differentiated samples) and six samples of VICs (including three under standard culture conditions and three differentiated samples), each with two technical replicates. The RNA-sequencing count analysis was performed in R using the DESeq2 library per the standard algorithm^17^. The VICs and osteoblast samples were compared at each time point, encompassing the standard culture conditions (control) and at the 10th day of osteogenic differentiation. The technical replicates were consolidated using the collapseReplicates function from the DESeq2 package. During data filtering, genes with counts below 10 were excluded. Statistical analysis involved the Wald test integrated into the standard DESeq2 analysis algorithm. Differentially expressed genes were chosen based on |Log2 Fold Change| > 1 and adjusted p-value < 0.05 criteria. Data normalization entailed applying the rlog transformation. The sample clustering was executed through Principal Component Analysis (PCA), and the conversion of Entrez Gene identifiers was conducted using the bitr function sourced from the clusterProfiler library.^18^

Enrichment analysis was performed using the clusterProfiler library, drawing from the Reactome Pathways database^19^ and human whole genome annotation org.Hs.eg.db. The differential gene expression results were visualized through various R packages, including pheatmap^20^, ComplexHeatmap^21^, EnhancedVolcano^21^, and ggplot2^22^. Additionally, Venn diagrams were generated using the VennDiagram library^23^ to illustrate the intersections among differentially expressed genes from VIC and osteoblast comparisons of varying types.

#### Proteomics data analysis

We performed data filtration and imputation of missing values with the NAguideR package for proteomic data analysis.^24^ Proteins with missed values in more than half of samples in at least one biological group and proteins with a coefficient of variation higher than 0.8 were removed. Then, we selected the optimal method for missed values imputation using “classic criteria” which was the “Robust data imputation” approach (“Impseqrob”).^25^

Differential expression analysis was performed similarly to RNA-seq data analysis described above, with minor differences. Specifically, we used quantile normalization instead of rlog transformation and limma instead of DESeq2 for proteomics data.^26^

## 3. Results

### 3.1 Various proteomics approaches give different proteomics coverage

To obtain maximal proteome coverage and estimate the effectiveness of various types of MS data acquisition on a timsTOF Pro platform for the real biological experiment, we collected three datasets for the comparison of osteogenic differentiation of valve interstitial cells (VICs) and osteoblasts: (1) classic data-dependent acquisition (DDA) in parallel accumulation serial fragmentation (PASEF) mode of ion mobility trapped mass-spectrometry, (2) data-independent acquisition (DIA) mass-spectrometry in PASEF mode with the identification of proteins based on empirically obtained spectral library or (3) with the identification based on “library-free” search with spectral library predicted from protein sequences by machine learning in DIA-NN software (DIA-ML).

After filtration, we included 2 563 proteins for DDA, 4 105 proteins for DIA, and 4 874 for the DIA-ML datasets for further analysis (supplementary materials 1). So, in our case, classic DIA gives 37.5% higher proteomics coverage than the DDA approach. Summarizing proteins, we found 5 405 proteins identified by at least one method and 2195 identified by all three. By RNA-seq data, we identified transcripts associated with 9941 genes, and it might be a biological reference – it is pretty unlikely for protein to be identified by shotgun proteomics data but not to be identified at mRNA level by RNA-seq. As expected, most proteins were identified by proteomics and RNA-seq data for classic DDA and DIA datasets. At the same time, 5% of all proteins identified by DIA-ML were unique for this approach and were not identified in RNA-seq data (fig. 2 a). This might be a sign of the weaker accuracy of this approach, which is expected.

**Figure 2.**
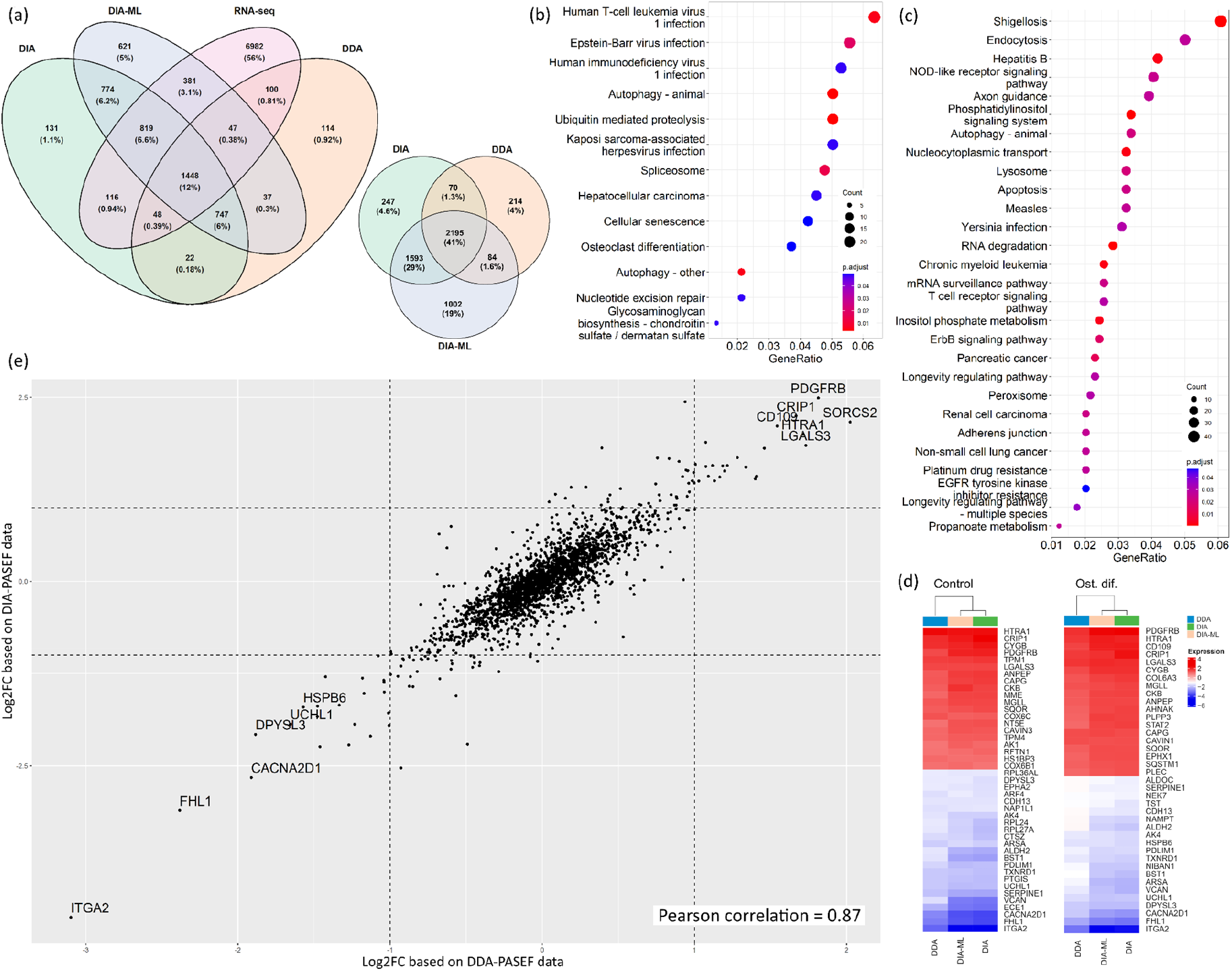
Comparison of three proteomics datasets (DDA, DIA, and DIA-ML) and RNA-seq data obtained for shotgun proteomics analysis of mechanisms of osteogenic differentiation of human valve interstitial cells (VICs) and osteoblasts. (a) Venn diagrams with overlap between gene products identified by RNA-seq in three proteomics datasets (left) and between gene products identified in three proteomics datasets (right). Proteomics datasets are represented after missed values filtration. (b, c) KEGG pathway enrichment analysis of proteins identified only by DIA-ML (b) or DIA (c). (d) Heatmap demonstrating the Log_2_ Fold Changes of differentially expressed genes, identified by all three proteomics methods between osteoblasts and differentiated VICs in standard cultivation (control) and after osteogenic differentiation (Ost. dif.). (e) Correlation of Log_2_ Fold Changes for differentially expressed proteins compared to differentiated osteoblasts and VICs by DDA and DIA proteomics.

Furthermore, all three proteomics datasets identified only 41% of proteins (fig. 2 b), 747 of which were not identified by RNA-seq (fig. 2 a). DIA should be able to identify more proteins, and we see 1593 proteins identified only in DIA and DIA-ML datasets. Nevertheless, we see 1002 proteins unique for DIA-ML data besides these DIA-unique proteins.

KEGG pathway enrichment analysis of proteins, unique for DIA-PASEF methods, demonstrates various functional groups, including several signaling pathways: ErbB, NOD-like receptor, T cell receptor, and phosphatidylinositol signaling system (fig. 2 c). Proteins, unique for DIA-ML data, are mainly associated with RNA and, therefore, with KEGG signaling pathways related to viral infections (fig. 2 d). Nonetheless, the term “Osteoclast differentiation” attracts attention in the context of osteogenic differentiation. In the DIA-ML dataset, this term includes several proteins important in the focus of our study: TAB2 (MAP3K7 binding protein 2), PIK3CB (phosphatidylinositol-4,5-bisphosphate 3-kinase catalytic subunit beta), PIK3R2 (phosphoinositide-3-kinase regulatory subunit 2), ITGB3 (integrin subunit beta 3), PPP3CB and PPP3CC (protein phosphatase 3 catalytic subunit beta and gamma), CTSK (cathepsin K), FOSL1 (FOS like 1, AP-1 transcription factor subunit), CHUK (a component of inhibitor of nuclear factor kappa B kinase complex), TRAF2 and TRAF6 (TNF receptor-associated factor 2 and 6).

Despite such differences in proteome coverage, these methods seem similar in the quantitation of proteins identified by both methods. We see a good correlation between different methods in pairwise Pearson correlation. The average correlation for the same sample in different datasets is 0.87 (0.83 – 0.88) between DDA and DIA; 0.87 (0.84 – 0.88) between DDA and DIA-ML; 0.98 (0.97 – 0.98) between DIA and DIA-ML. Accordingly, there is a good correlation between Log_2_ Fold Changes calculated in different datasets for pairwise comparisons of biological groups studied (see detailed discussion further). The average correlation Log_2_ Fold Changes in different datasets is 0.83 (0.8 – 0.87) between DDA and DIA; 0.83 (0.81 – 0.86) between DDA and DIA-ML; 0.91 (0.89 – 0.92) between DIA and DIA-ML. We want to emphasize the apparent correlation between Log2 Fold Changes for differentially expressed proteins and the complete absence of proteins with mutually exclusive expression in different datasets (fig. 2 e, supplementary materials 2).

### 3.2 Proteomic and transcriptomic profiles of valve interstitial cells are distinct from osteoblasts both before and after osteogenic differentiation

Using the four obtained datasets, we compared the osteogenic differentiation of valve interstitial cells (pathological calcification) and osteoblasts (normal ossification). Clustering of the samples by principal component analysis (PCA) revealed a similar pattern for all datasets (supplementary materials 3) – VICs form distinct clusters before and after osteogenic differentiation and never cluster together with osteoblasts (fig. 3 a). Similar to our previous studies, osteoblasts form mixed clusters (fig. 3 a).^27^

**Figure 3.**
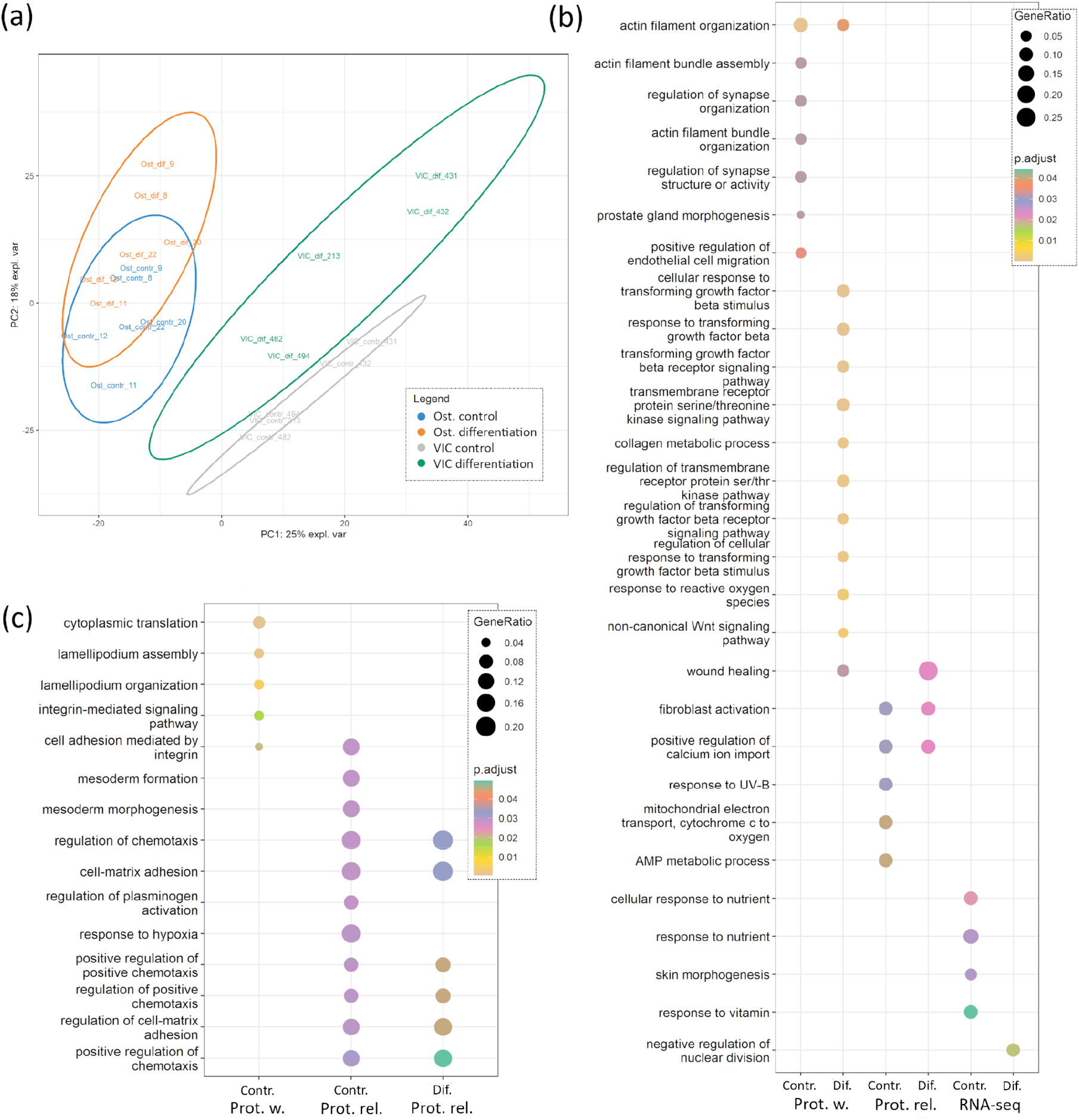
Comparison of human valve interstitial cells (VICs) and osteoblasts (Ost) cultured in standard conditions or after induction of osteogenic differentiation. (a) Clusterization of the samples in the principal component analysis (PCA) dimension based on DIA-PASEF data. Gray – VICs in standard cultivation; Green – VICs on the 10^th^ day after induction of osteogenic differentiation; Blue – osteoblasts in standard cultivation; Yellow – osteoblasts on the 10^th^ day after induction of osteogenic differentiation. (b) Proteins or transcripts upregulated in osteoblasts during standard cultivation (Contr.) or on the 10^th^ day after induction of osteogenic differentiation (Dif.). (c) Proteins or transcripts upregulated in VICs during standard cultivation (contr.) or on the 10^th^ day after induction of osteogenic differentiation (Dif.). RNA-seq – transcripts identified by RNA-seq; Prot. w. – differentially expressed proteins, identified by at least one shotgun proteomics dataset; Prot. rel. – differentially expressed proteins identified by all three shotgun proteomics datasets.

For optimal data representation, we combined the results of the analysis of a differential expression to the three datasets: (1) differentially expressed transcripts identified by RNA-seq, (2) differentially expressed proteins identified in all three proteomics datasets (Proteomics reliable, “Prot. rel.”), or (3) identified in at least one of the proteomics datasets (Proteomics wide, “Prot. w.”). The datasets include 101, 20, and 162 genes upregulated in osteoblasts; 106, 23, and 208 genes upregulated in VICs, respectively, in standard cultivation. 102, 19, and 206 genes were upregulated in osteoblasts; 118, 20, and 191 genes were upregulated in VICs, respectively, after the induction of osteogenic differentiation.

In focusing on the gene ontology analysis in “Biological Processes”, a limited intersection of differentially expressed genes is noticeable before and after the induction of osteogenic differentiation (fig. 3). Noteworthy is the Prot. rel dataset encompasses proteins identified via all three proteomic methods (DIA, DDA, and DIA-ML). In osteoblasts, this dataset shows overlaps in the biological processes of fibroblast activation and positive regulation of calcium ion import. In VICs, a more diverse array of processes is observed, primarily associated with chemotaxis, cell adhesion to the extracellular matrix, and regulating these processes.

Under standard culture conditions (control), osteoblasts exhibit the activation of actin filament assembly and organization processes, fibroblast activation, and cellular response to nutrients. In VICs under the same conditions, genes are enriched in lamellipodium assembly and organization, mesoderm formation, cell-matrix adhesion, and the positive regulation of chemotaxis. Differentiated osteoblasts show an upregulation of TGF-beta signaling and non-canonical Wnt signaling, an increase in collagen metabolism, and the activation of fibroblasts (fig. 3b). In differentiated VICs, these processes are absent, and the enriched pathways coincide with the pathways identified in undifferentiated VICs (fig. 3c). It is noteworthy that only about 30% of differentially expressed genes significantly differ between both control and differentiated VICs and osteoblasts. At the same time, about 60% of the genes are differentially expressed between these cell types only before or after osteogenic differentiation (supplementary materials 3).

We found a low overlap between proteomics and RNA-seq data, which was expected from our previous proteotranscriptomics dataset of VICs osteogenic differentiation (fig. 4).^28^ Table 2 lists gene products (proteins or transcripts) up-or downregulated between osteoblasts and VICs in both proteomics and RNA-seq data.

**Table 2.** Genes, products of which are up-or downregulated in both RNA-seq and one of the proteomic datasets in comparisons of osteoblasts and valve interstitial cells (VICs) in standard cultivation conditions or on the 10^th^ day of osteogenic differentiation.

|  | Standard cultivation | Osteogenic differentiation |
| --- | --- | --- |
| Upregulated in osteoblasts | **TGFBI, SULF1, GALNT1, CALD1, POSTN** | **TGFBI, SERPINE2, COL8A1** |
| Upregulated in VICs | **VCAN, EFEMP1, ANK3, CCDC80, CACNA2D1** | **VCAN, EFEMP1, ANK3, CCDC80, HEG1** |

One of the differences between transcriptomic and proteomic datasets is that in the RNA-seq data, we found a surprisingly high proportion of differentially expressed long non-coding RNA (lncRNA). Most of these lncRNAs are known antisense RNA (tab. 3). Notably, many of these antisense RNAs significantly differ only before but not after induction of osteogenic differentiation (tab. 3).

**Table 3.** Annotated antisense RNA found in transcriptomes of osteoblasts in comparison with valve interstitial in standard cultivation conditions or on the 10^th^ day of osteogenic differentiation. Non-significant – adjusted P-value > 0.05

| Upregulated in osteoblasts |  |  |  |  |  |  |
| --- | --- | --- | --- | --- | --- | --- |
| Name | ENSEMBL | Std. cultivation |  | Ost. differentiation |  | Target gene |
|  |  | Log<sub>2</sub> FC | adj. p.val. | Log<sub>2</sub> FC | adj. p.val. |  |
| HOXA-AS3 | ENSG00000254369 | 8.3 | 4.82E-09 | 7.7 | 2.02E-07 | HOXA |
| HOXA10-AS | ENSG00000253187 | 8.2 | 9.84E-09 | 7.1 | 1.71E-06 | HOXA10 |
| DLX6-AS1 | ENSG00000231764 | 6.0 | 2.76E-04 | 7.0 | 1.86E-05 | DLX6 |
| HOXA11-AS | ENSG00000240990 | 6.8 | 2.38E-05 | 5.9 | 1.93E-03 | HOXA11 |
| Lnc-FIGNL2-6 | ENSG00000260122 | 5.8 | 7.42E-04 | 5.7 | 1.06E-03 | HSALNG0091106 |
| CHL1-AS1 | ENSG00000234661 | 6.2 | 3.47E-04 | 5.6 | 2.38E-03 | CHL1 |
| Lnc-RD3-11 | ENSG00000279333 | 7.6 | 5.50E-07 | 5.6 | 1.32E-03 | KCNH1 |
| PENK-AS1 | ENSG00000254254 | 5.2 | 4.38E-04 | 5.4 | 9.61E-08 | PENK |
| HOXD-AS2 | ENSG00000237380 | non-significant |  | 5.2 | 2.55E-02 | HOXD |
| Lnc-DAOA-47 | ENSG00000284966 | 4.1 | 2.01E-10 | 5.1 | 7.12E-02 | EFNB2 |
| Lnc-SLC25A32-9 | ENSG00000253851 | 6.5 | 6.51E-07 | non-significant |  | BAALC |
| Upregulated in valve interstitial cells |  |  |  |  |  |  |
| Name | ENSEMBL | Standard cultivation |  | Osteogenic differentiation |  | Target gene |
|  |  | Log<sub>2</sub> FC | adj. p.val. | Log<sub>2</sub> FC | adj. p.val. |  |
| Lnc-COPG2-3 | ENSG00000270823 | 1.8 | 5.13E-07 | 6.4 | 2.24E-05 | MEST |
| LAMA5-AS1 | ENSG00000228812 | non-significant |  | 5.1 | 8.59E-03 | LAMA5 |
| CAVIN2-AS1 | ENSG00000233766 | 3.5 | 3.94E-02 | 5.0 | 1.08E-02 | CAVIN2 |
| ANK3-AS | ENSG00000232682 | 3.6 | 1.73E-19 | 4.9 | 3.14E-37 | ANK3 |
| CADM3-AS1 | ENSG00000225670 | 6.6 | 8.84E-03 | non-significant |  | CADM3 |
| CCDC110-AS | ENSG00000249679 | 6.07 | 1.34E-04 | 4.3 | 3.23E-02 | CCDC110 |
| ADGRD1-AS1 | ENSG00000256151 | 6.0 | 5.02E-04 | non-significant |  | ADGRD1 |
| Lnc-THNSL1-4 | ENSG00000273107 | 5.9 | 9.12E-04 | non-significant |  | THNSL1 |
| HAND2-AS1 | ENSG00000237125 | 5.1 | 5.05E-09 | 2.5 | 1.36E-04 | HAND2 |
| ENSG00000289061 | ENSG00000289061 | 5.1 | 1.84E-02 | non-significant |  | EMILIN2 |

## 4. Discussion

### 4.1 DIA-PASEF with library-free search is optimal for routine shotgun proteomics

The rapid advancements in shotgun proteomic techniques present a challenge for end users, particularly those who may need to be more experts in proteomics, to determine the most suitable approach for their specific needs. Furthermore, multiple data acquisition modes are available within a single proteomics platform, making it challenging to select the optimal one.

Trapped Ion Mobility Spectrometry (TIMS), utilized in the timsTOF Pro platform, stands out as one of the leading technologies for routine shotgun proteomics. It is particularly valuable for analyses involving limited sample amounts or the need to study large cohorts, as often found in biomedical research. However, comprehensive comparisons of different data acquisition and analysis strategies are scarce from a user’s perspective, particularly those conducted within actual biological objectives and complemented by transcriptomics data.

In this context, we have conducted a comparative study of two major proteomic approaches on the timsTOF Pro platform: Data Dependent Acquisition (DDA) and Data Independent Acquisition (DIA) with Parallel Accumulation-Serial Fragmentation (PASEF). We used one of the most modern approaches for data analysis, but it is still available in a user-friendly interface. We used user-friendly approaches: MSFragger for DDA-PASEF and DIA-NN for DIA-PASEF data. ^29,30^

Our biological aim was to identify potential therapeutic targets for the selective inhibition of osteogenic differentiation in valve interstitial cells (VICs), an essential process in CAVD. This comparison sheds light on the efficacy of these approaches for specific applications and bridges the gap between proteomic and transcriptomic analyses in searching for effective CAVD treatments.

From a technical standpoint, our samples exhibit greater sophistication than the HeLa samples typically utilized in comparative proteomics research. This sophistication arises from three key factors: (i) The primary function of both VICs and osteoblasts is the secretion of extracellular matrix (ECM). As a result, their proteomes are enriched with numerous ECM proteins, which interfere with identifying less abundant proteins, thereby limiting proteome coverage. (ii) The primary cell cultures in our study are derived from donors with diverse genetic backgrounds and lifestyle factors, introducing a more comprehensive range of biological variability compared to studies using uniform cell lines like HeLa. (iii) The availability of a limited number of biological replicates is a common constraint in most biomedical studies, including ours. This limitation can affect the statistical power and reproducibility of the findings.

As a result, our study quantified half as many protein groups as those reported for HeLa cells.^13^ However, DIA-PASEF methodologies proved to be more advantageous in our context. Specifically, using DIA-NN library-free search and the conventional library-based DIA analysis increased the quantification of protein groups to 90% and 60%, respectively. A similar dramatic increase in performance was observed in the comparison of DDA-and DIA-PASEF methods for metaproteomics, which possess even more technical challenges than we faced in our study. In the study of Gómez-Varela et al., DIA-PASEF resulted in the quantification of up to 4 times more taxonomic units than DDA-PASEF.^12^ This contrasts with the findings of Huang et al., who reported a mere 30% increase in quantified proteins for HeLa samples using a similar DDA-and DIA-PASEF comparative analysis.

To evaluate the reliability of our protein identification, we compared the identified proteins with the transcripts detected through RNA-seq transcriptomics. Given the inherent discrepancies expected between RNA-seq and proteomics data, especially in human primary cell lines undergoing differentiation^28^, this comparative analysis is limited. Still, it might be used to assess various PASEF techniques. Notably, the highest number of unique proteins was observed with DIA-PASEF data analyzed through a library-free approach, accounting for only 5% of all proteins identified (Fig. 2a). Meanwhile, traditional DIA-PASEF using a library-based search yielded the most balanced results in terms of both high protein quantification and methodological overlap. This finding aligns well with existing literature recognizing library-based searches as exceptionally reliable for analyzing DDA and DIA data.^31^

Achieving nearly a twofold increase in the number of proteins identified is critical for the downstream identification of differentially expressed proteins. This significance is underscored by the fact that highly abundant proteins are often housekeeping proteins, which are remarkably stable from both evolutionary and functional perspectives.^32,33^ In our study, this enhanced identification capability has enabled us to detect hundreds of proteins associated with key cellular compartments (such as peroxisomes and lysosomes) and processes (including endocytosis and apoptosis), as well as numerous signaling pathways, such as ErbB, NOD-like receptor, phosphatidylinositol, T-cell receptor, and EGFR (Fig. 2b-c). As a result, we significantly increased the detection of differentially expressed proteins (DEPs), identifying 371 DEPs in DDA datasets and 646 DEPs in DIA datasets when comparing valve interstitial cells (VICs) and osteoblasts after osteogenic differentiation as an example (Supplementary Materials 1).

Further in-depth comparison of DEPs identified by different PASEF protocols revealed a surprisingly high correlation in DEP identification across various methods. The correlation of Log2 Fold Changes (Log2FC) between different approaches within all biological comparisons ranged from 0.8 to 0.92. We encountered no DEPs with opposing directions across the datasets (Fig. 2d-e, Supplementary Materials 2). The highest correlation was observed between DIA and DIA-ML approaches (0.91), indicating a robust consistency in our data.

In summary, our observations have shown that the DIA-PASEF method offers a significantly greater advantage over the DDA approach for complex samples in actual proteomics studies, surpassing the performance observed with HeLa test samples. While the library-based search in DIA-PASEF sets a high standard in shotgun proteomics, the DIA-NN-based library-free search presents a more accessible alternative. The library-free search, facilitated by the user-friendly DIA-NN software, rivals the library-based approach in terms of differentially expressed proteins (DEPs) identified, and it only exhibits a slight reduction in data reliability. Our findings suggest that for routine shotgun proteomics tasks on the timsTOF Pro platform, DIA-PASEF with a library-free search emerges as the preferable method.

### 4.2 Specificity of VICs osteogenic differentiation offers potential for target CAVD therapy

The comprehensive multi-omics dataset discussed in detail above allows us to investigate the differences in molecular mechanisms of normal osteogenic differentiation in osteoblasts and pathological osteogenic differentiation in valvular interstitial cells (VICs).

Calcified aortic valves, particularly in the later stages of calcific aortic valve disease (CAVD), are often presumed to exhibit a high degree of similarity to lamellar bone regarding calcium crystal structure and histological organization. However, upon closer examination, we can identify fundamental differences even at the histological level. Mohler et al. (2001) conducted a study on heterotopic ossification in 347 surgically removed heart valves and discovered that while microcalcification was present in most of the cases, mature lamellar bone with hematopoietic elements was observed only in 13% of the cases, with signs of endochondral bone formation being rare and identified in 4 valves. Most valves undergoing ossification exhibited microfractures and neoangiogenesis. This suggests that structures akin to mature bone are only observed in a minority of cases. Predominantly, microcalcification is observed alongside microfractures and inflammation, which significantly diverges from the processes involved in normal ossification.^34^

Consequently, our observations reveal significant differences in the mechanisms of osteogenic differentiation between osteoblast-like cells derived from adult bone and VICs isolated from calcified aortic valves in the CAVD latter stages. As previously demonstrated, the osteoblast proteome does not undergo significant changes during osteogenic differentiation.^27,35^ In contrast, the VICs’ proteome exhibits notable alterations (Supplementary Materials 3).^28^ Before and following differentiation, VICs express higher levels of proteins related to chemotaxis, cell-matrix adhesion, and mesodermal markers. Conversely, osteoblasts show an increased expression of proteins involved in osteogenic differentiation, such as those associated with TGF-beta and Wnt signaling pathways, collagen synthesis, and calcium ion import. Given the endothelial origins of VICs^7^, these differences align well with previous studies that have documented varied osteogenic differentiation mechanisms in cells of different origins. ^9,35–39^ Furthermore, research by Nantavisai et al. (2020) suggests that these distinctions could be leveraged for the selective inhibition of osteogenic differentiation in specific cell types *in vitro*.^36^ From this perspective, proteins exclusively upregulated during VICs’ osteogenic differentiation present promising targets for selective anti-CAVD treatments (supplementary materials 5).

Our findings reveal a significant overexpression of Monoamine oxidase A (MAOA) during osteogenic differentiation of valvular interstitial cells (VICs) compared to osteoblasts. Recognized as a critical marker for CAVD, MAOA has been identified as a potential treatment for this disease.^40^ Importantly, our data highlight its selectivity in targeting pathological valve calcification, enhancing its therapeutic promise. Additionally, we observed an upregulation of A-Kinase Anchoring Protein 2 (AKAP2) exclusively during VICs’ osteogenic differentiation. Previous studies have linked AKAP2 mutations with adolescent idiopathic scoliosis and demonstrated the critical role of the AKAP2/ERK1/2 signaling axis in longitudinal bone growth.^41,42^ The specific upregulation of AKAP2 in VICs—unlike in osteoblasts—suggests its association with the early stages of osteogenic differentiation in VICs. Similarly, another protein upregulated in VICs differentiation, GLIPR2, also modulates ERK1/2 signaling, though its role in osteogenic differentiation has not been previously described.^43^ The p38 mitogen-activated protein kinase (MAPK) signaling pathway, which includes ERK1/2, is involved in inflammation and is well-known to be associated with atherosclerotic and aortic valve calcification.^44^ Moreover, it was shown that andrographolide inhibits VICs calcification via the NF-kappa B/Akt/ERK pathway *in vitro*, which supports our data regarding ERK1/2 to be a promising target for selective therapy against aortic valve calcification.^45^

## Supplementary Materials

Supplementary Materials 1: Results of differential expression analysis between VICs and osteoblasts in control and osteogenic differentiation for the three proteomics and RNA-seq datasets; Supplementary Materials 2: Correlation of Log2 Fold changes for differentially expressed proteins between proteomic datasets; Supplementary Materials 3: Clustering of osteoblasts and human valve interstitial cells (VICs) samples by principal component analysis (PCA); Supplementary Materials 4: Venn diagrams representing the overlap of differentially expressed genes and proteins; Supplementary Materials 5: Boxplots for selective proteins

## Acknowledgment

Shotgun proteomics analysis was carried out using the equipment of the Core Facility “Centre for Molecular and Cell Technologies” (St. Petersburg State University). RNA sequencing was performed on the Core Facility “Medical Genomics” (Tomsk NRMC) equipment and the TomskRegional Common Use Center.

