## Supplemental Materials 2-5 for "Similar, but not the same: multi-omics comparison of human valve interstitial cells and osteoblast osteogenic differentiation expanded with an estimation of data-dependent and data-independent PASEF"

**This Supplementary file includes:**

Supplementary Materials 1: Results of differential expression analysis between VICs and osteoblasts in control and osteogenic differentiation for the three proteomics and RNA-seq datasets (separate xlsx file)

Supplementary Materials 2: Correlation of Log2 Fold changes for differentially expressed proteins between proteomic datasets

Supplementary Materials 3: Clustering of osteoblasts and human valve interstitial cells (VICs) samples by principal component analysis (PCA)

Supplementary Materials 4: Venn diagrams representing the overlap of differentially expressed genes and proteins

Supplementary Materials 5: Boxplots for selective proteins

**Supplementary Materials 2**. Correlation of Log2 Fold changes for differentially expressed proteins between proteomic datasets.

**Table 1**. Pearson correlation of Log2 Fold changes for differentially expressed proteins between DIA and DDA, DDA and DIA-ML, DIA and DIA-ML proteomes. Ost – correlation of Log2 Fold changes between differentiated and control (undifferentiated) osteoblasts. VIC – correlation of Log2 Fold changes between differentiated and control VICs (Valve Interstitial Cells). Ost VIC contr – correlation of Log2 Fold changes between control osteoblasts and control VICs. Ost VIC dif – correlation of Log2 Fold changes between differentiated osteoblasts and differentiated VICs.

| **Pearson correlation** | Ost | VICs | Ost VICs contr | Ost VICs dif |
|---|---|---|---|---|
| DDA vs DIA | 0.8310 | 0.8038 | 0.8210 | 0.8681 |
| DDA vs DIA-ML | 0.8257 | 0.8086 | 0.8198 | 0.8632 |
| DIA vs DIA-ML | 0.8894 | 0.8985 | 0.9157 | 0.9241 |

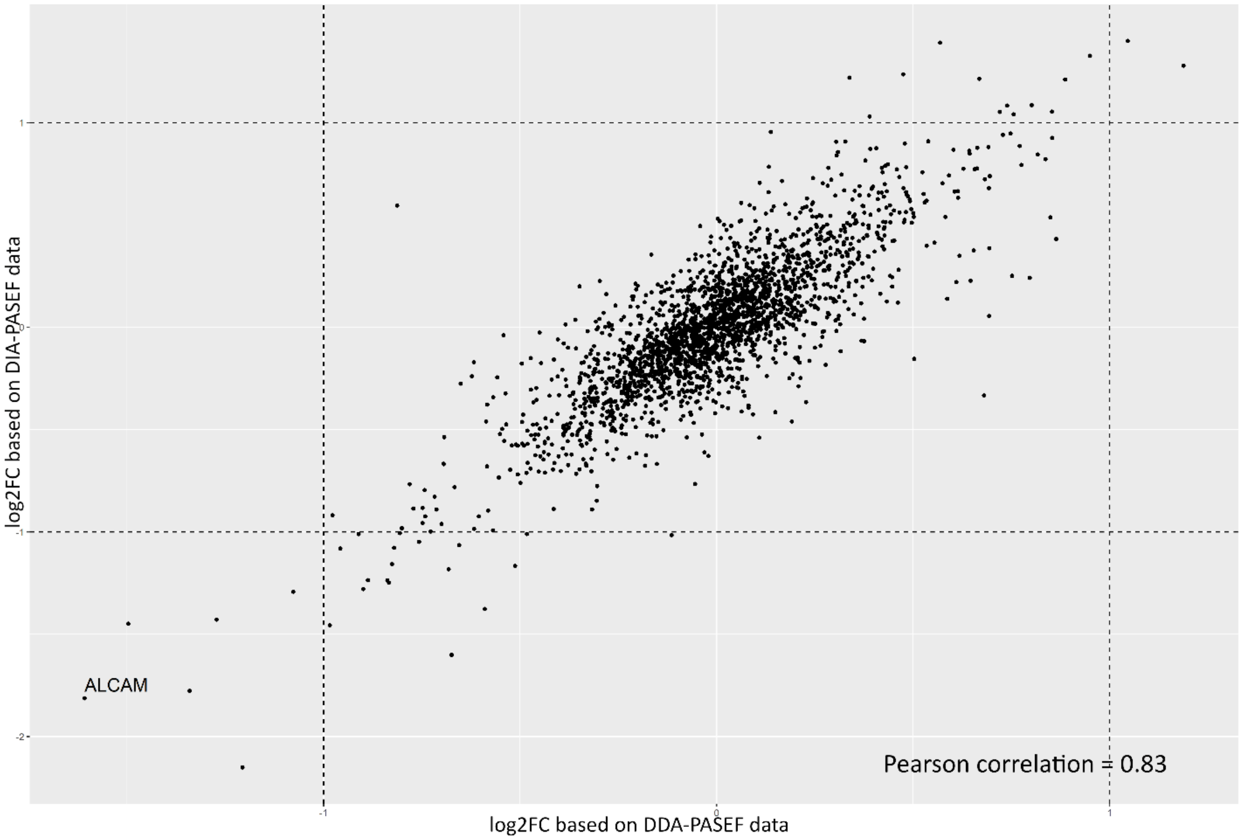


**Figure 1.** Correlation of Log2 Fold Changes for differentially expressed proteins in comparison of differentiated osteoblasts and control osteoblasts by DDA and DIA proteomics.

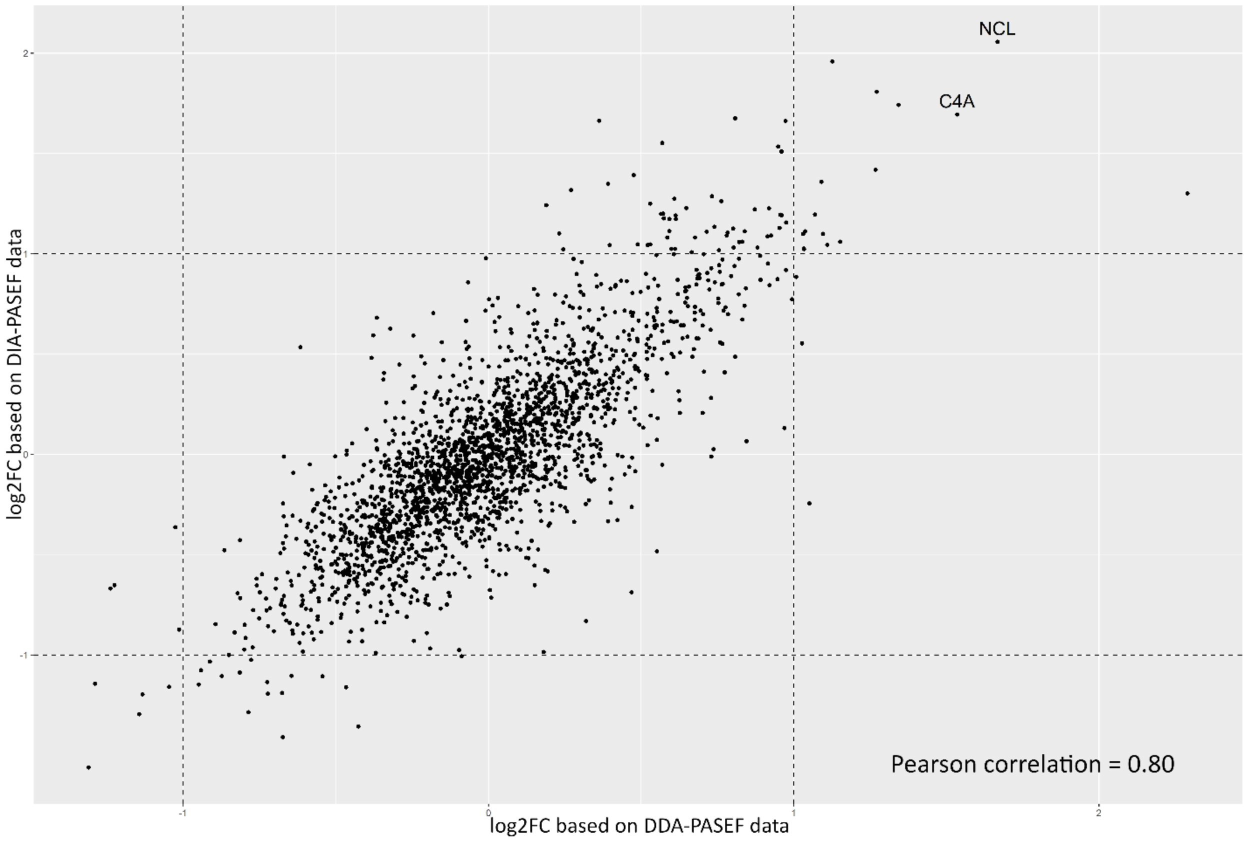


**Figure 2.** Correlation of Log2 Fold Changes for differentially expressed proteins in comparison of differentiated VIC and control VIC by DDA and DIA proteomics.

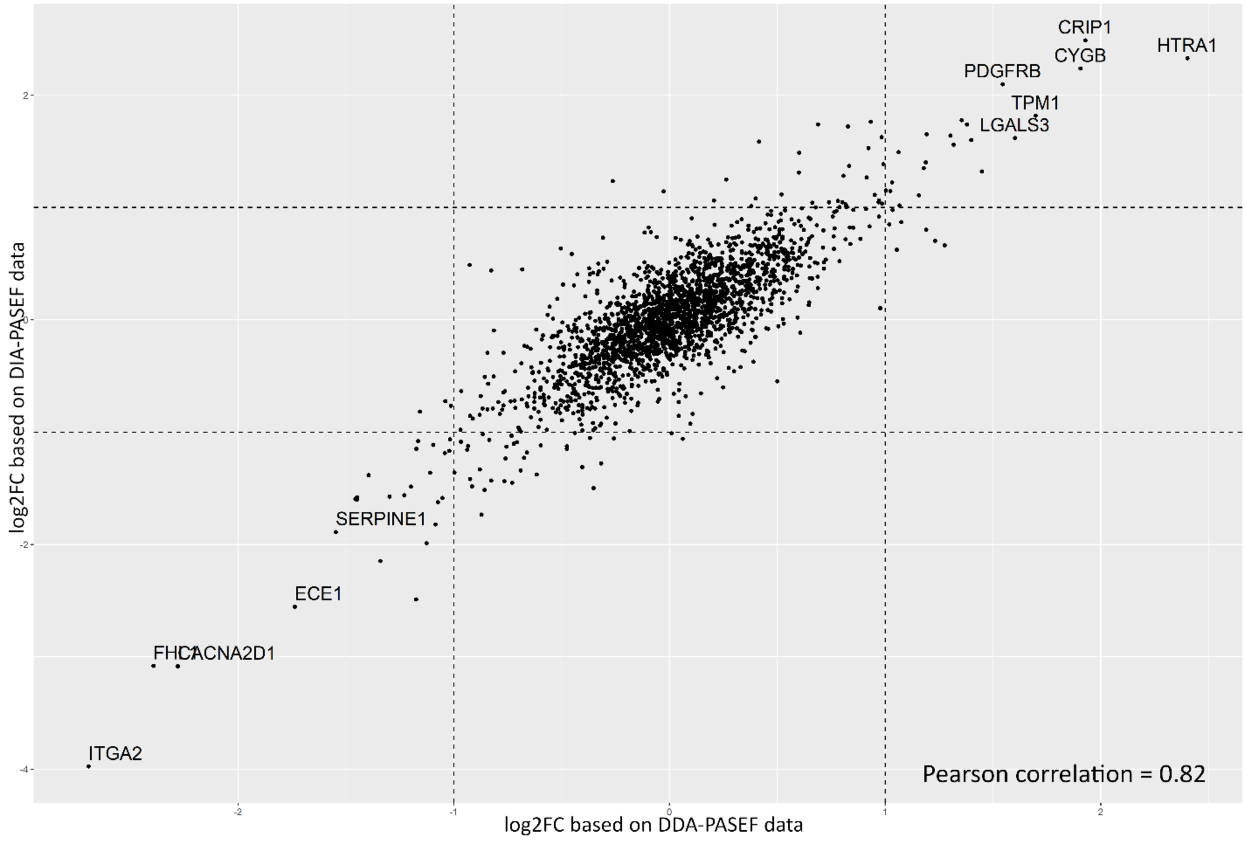


**Figure 3.** Correlation of Log2 Fold Changes for differentially expressed proteins in comparison of control osteoblasts and control VIC by DDA and DIA proteomics.

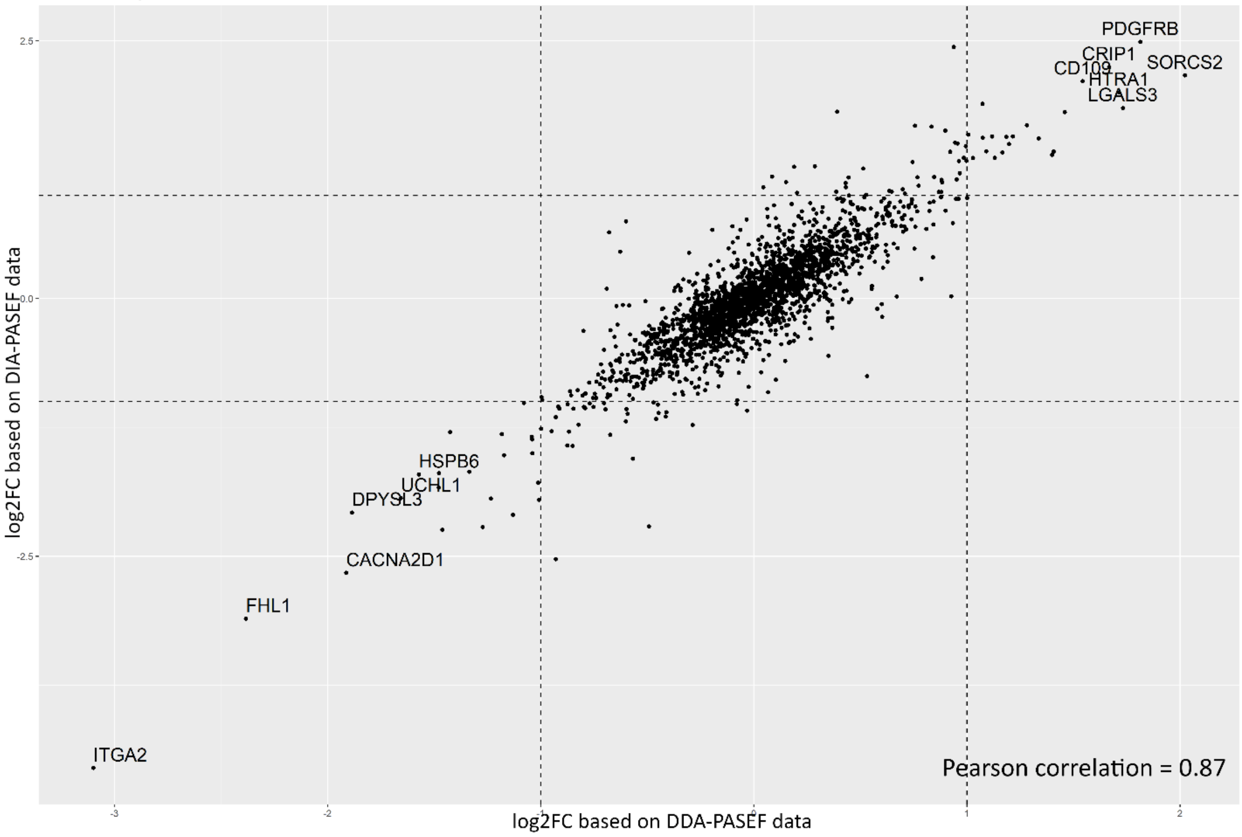


**Figure 4.** Correlation of Log2 Fold Changes for differentially expressed proteins in comparison of differentiated osteoblasts and differentiated VIC by DDA and DIA proteomics.

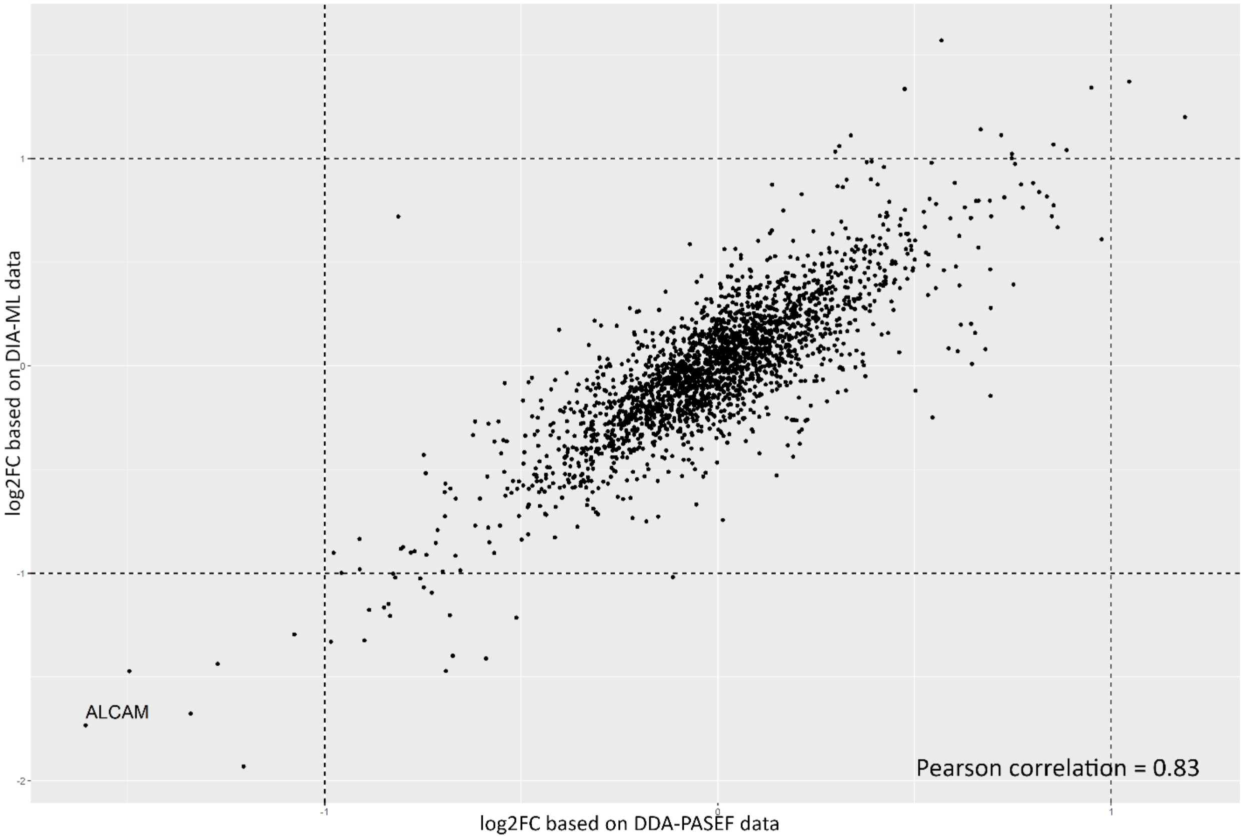


**Figure 5.** Correlation of Log2 Fold Changes for differentially expressed proteins in comparison of differentiated osteoblasts and control osteoblasts by DDA and DIA-ML proteomics.

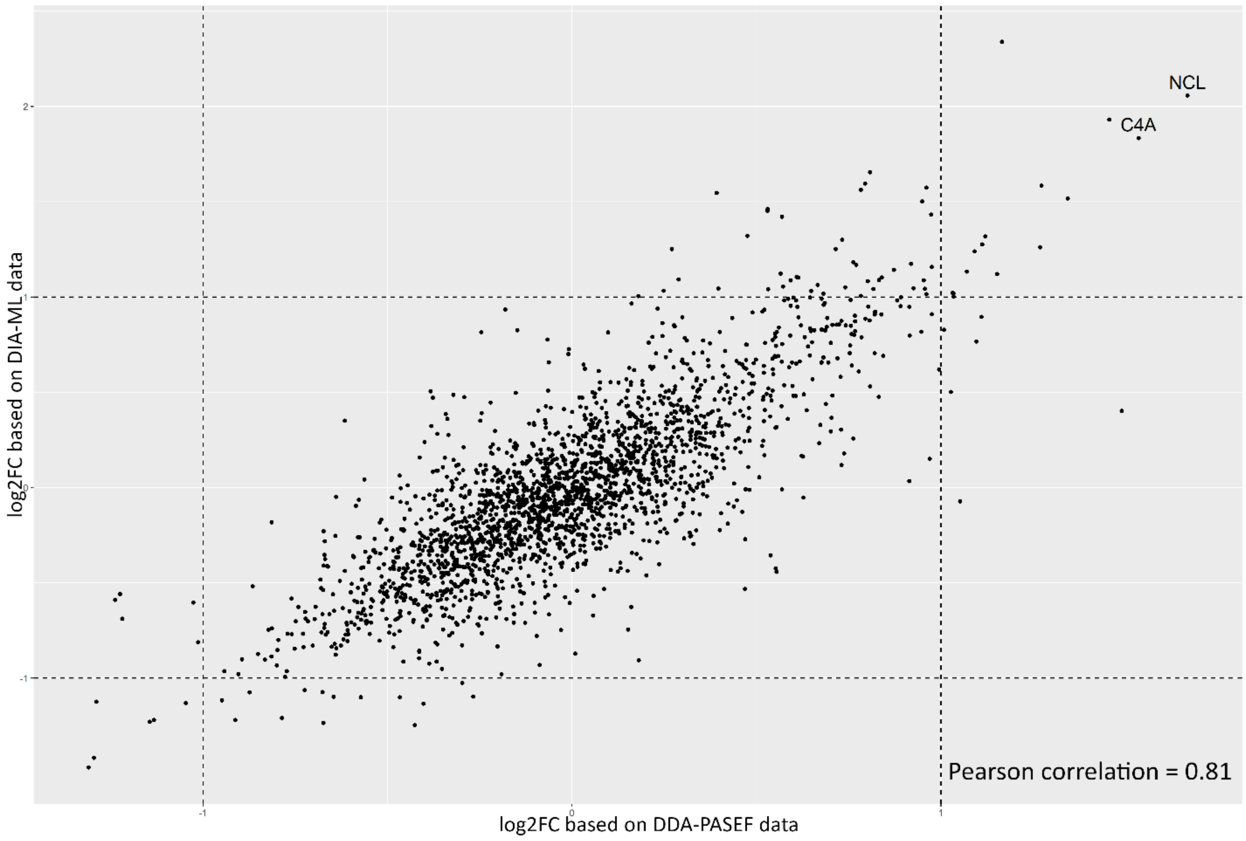


**Figure 6.** Correlation of Log2 Fold Changes for differentially expressed proteins in comparison of differentiated VICs and control VICs by DDA and DIA-ML proteomics.

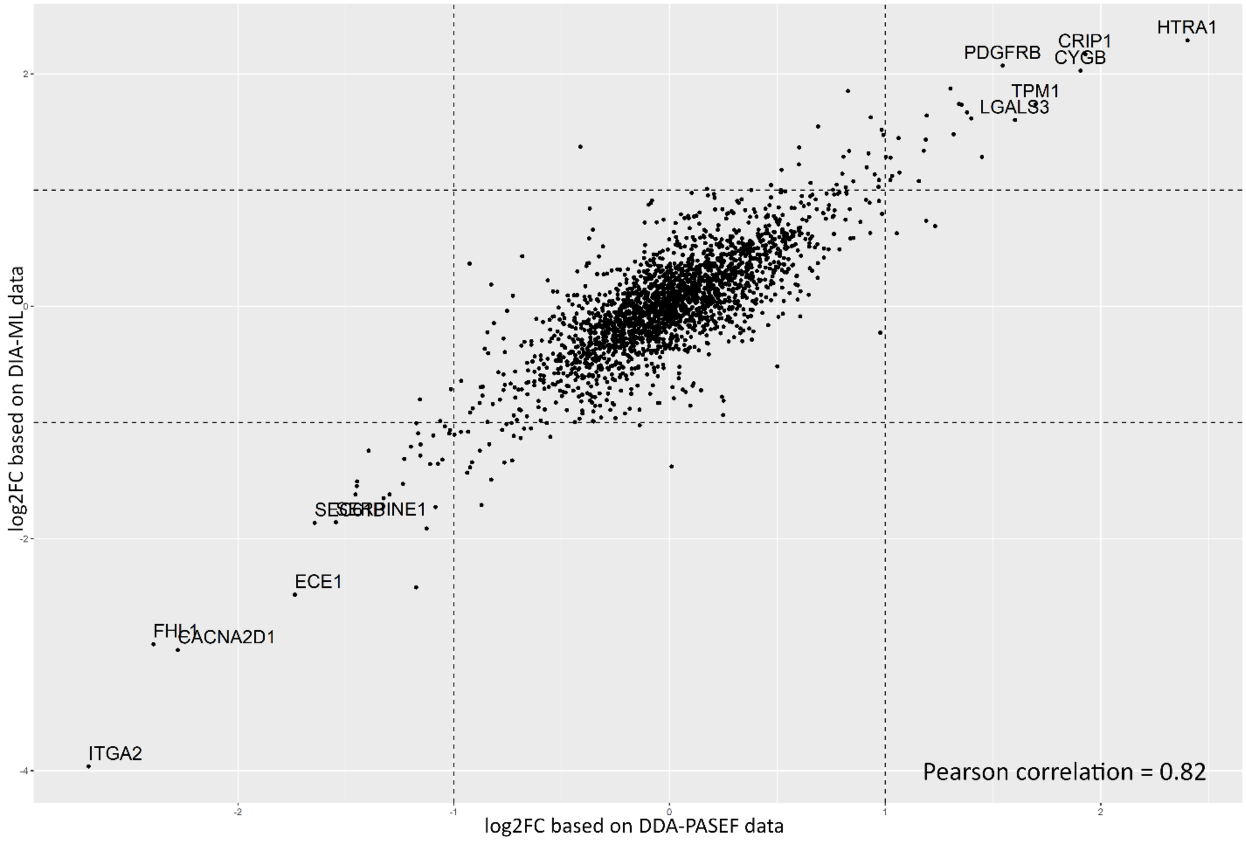


**Figure 7.** Correlation of Log2 Fold Changes for differentially expressed proteins in comparison of control osteoblasts and control VICs by DDA and DIA-ML proteomics.

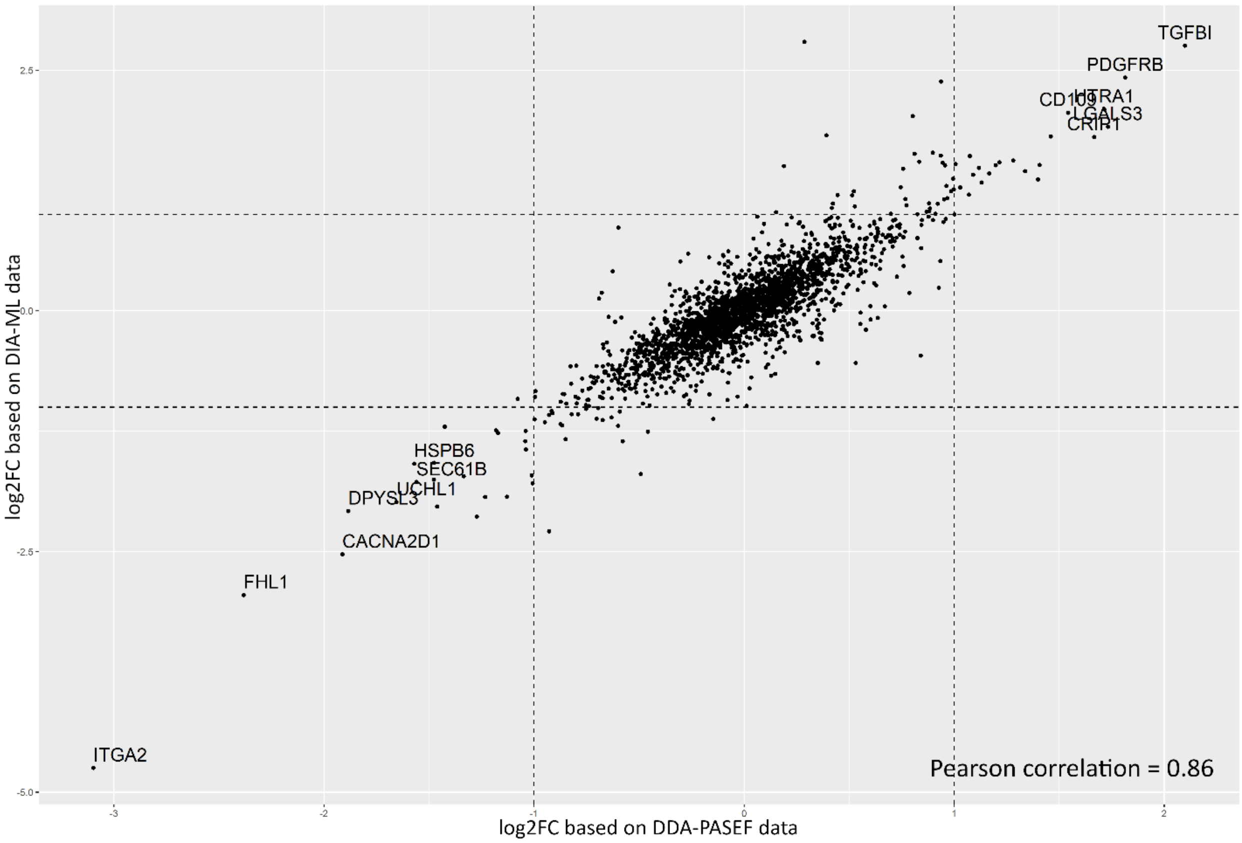


**Figure 8.** Correlation of Log2 Fold Changes for differentially expressed proteins in comparison of differentiated osteoblasts and differentiated VICs by DDA and DIA-ML proteomics.

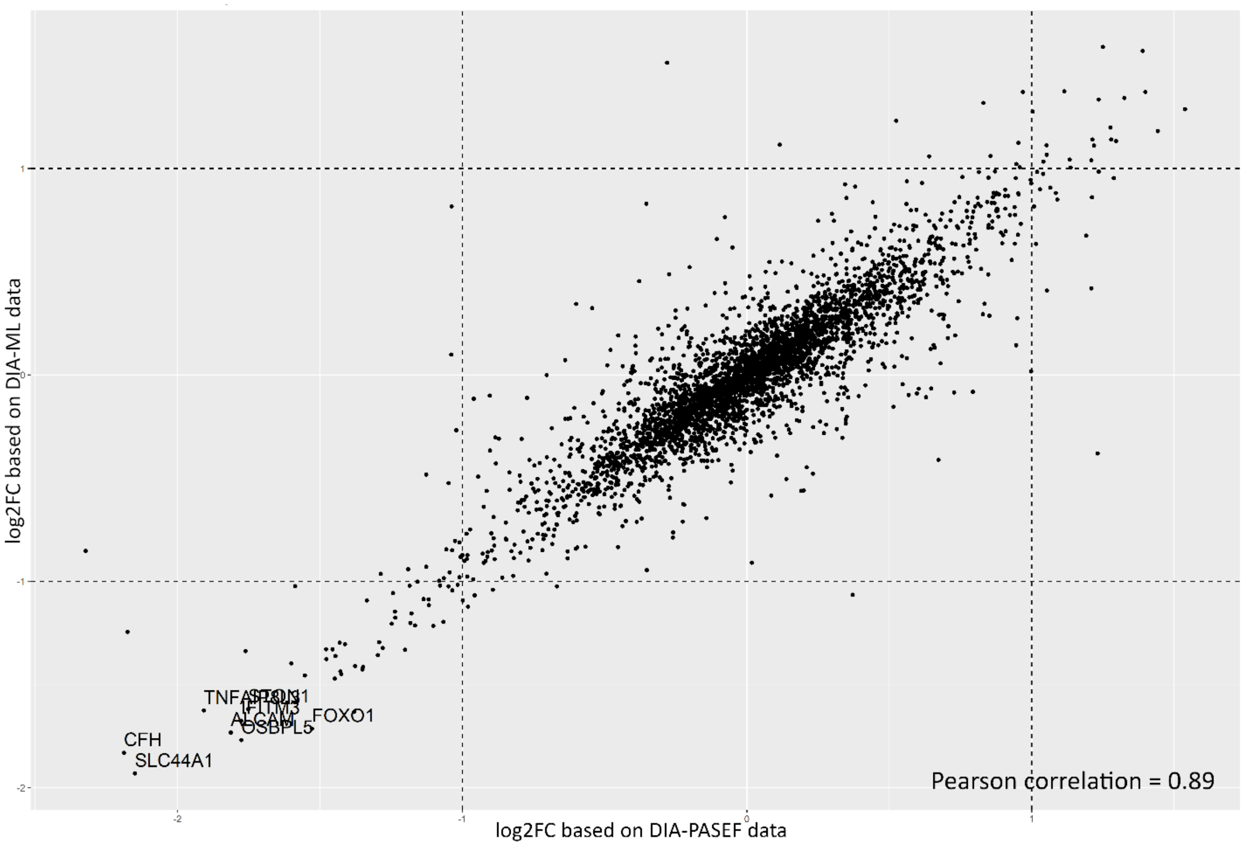


**Figure 9.** Correlation of Log2 Fold Changes for differentially expressed proteins in comparison of differentiated osteoblasts and control osteoblasts by DIA and DIA-ML proteomics.

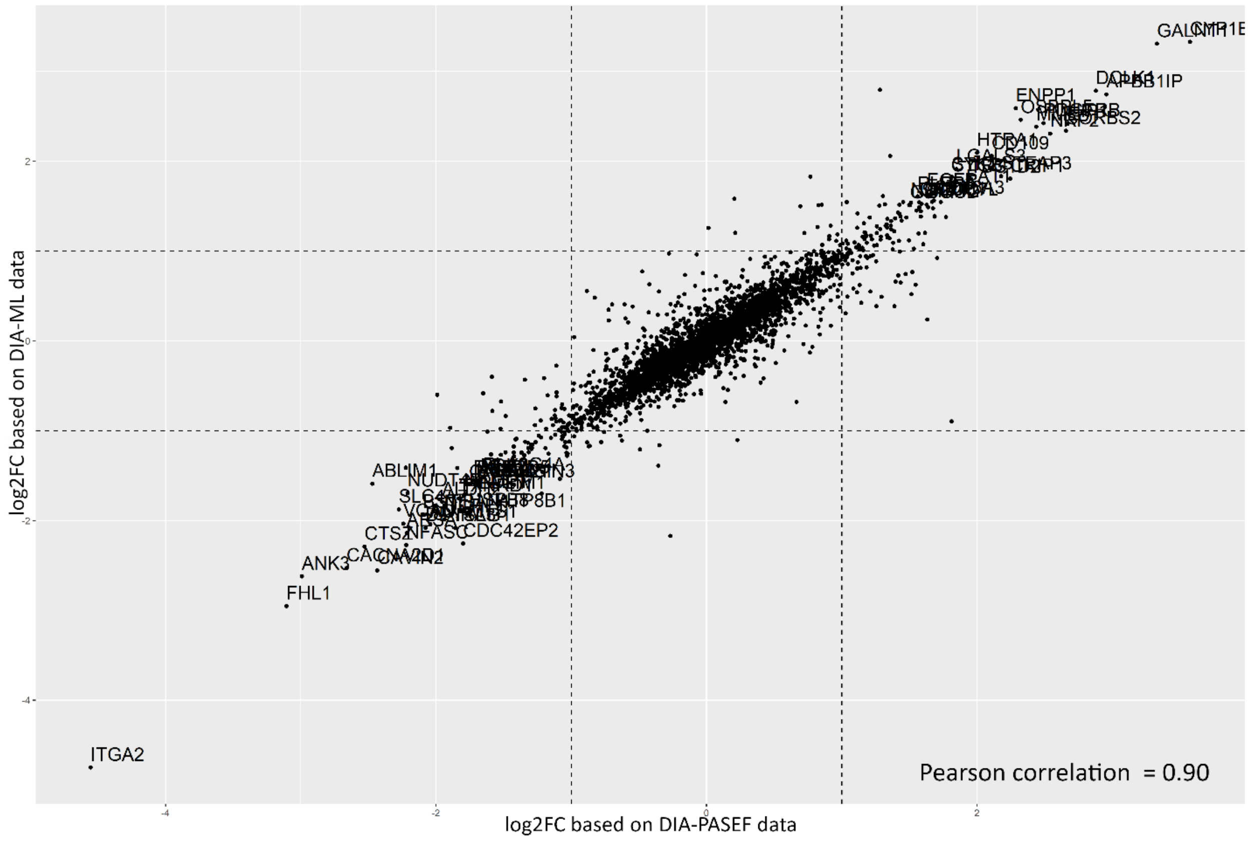


**Figure 10.** Correlation of Log2 Fold Changes for differentially expressed proteins in comparison of differentiated VICs and control VICs by DIA and DIA-ML proteomics.

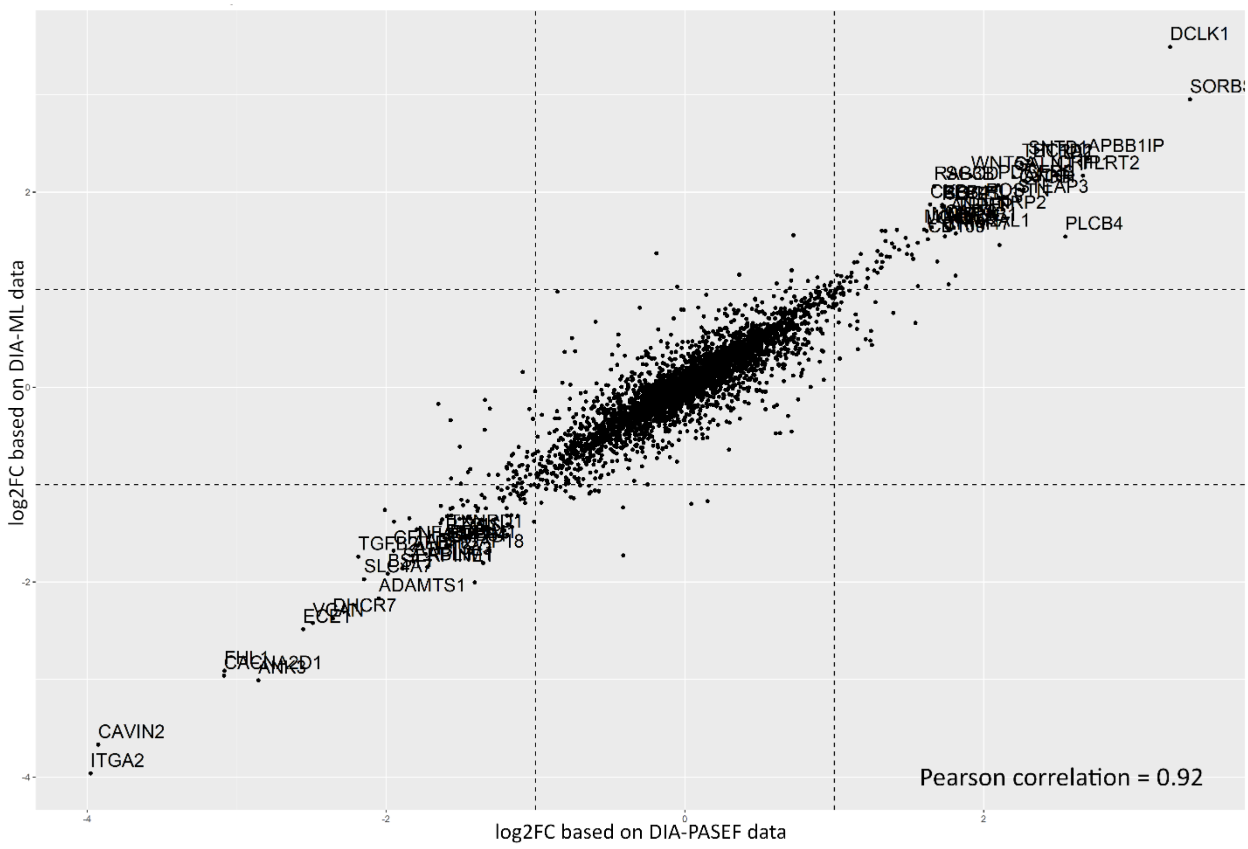


**Figure 11.** Correlation of Log2 Fold Changes for differentially expressed proteins in comparison of control osteoblasts and control VICs by DIA and DIA-ML proteomics.

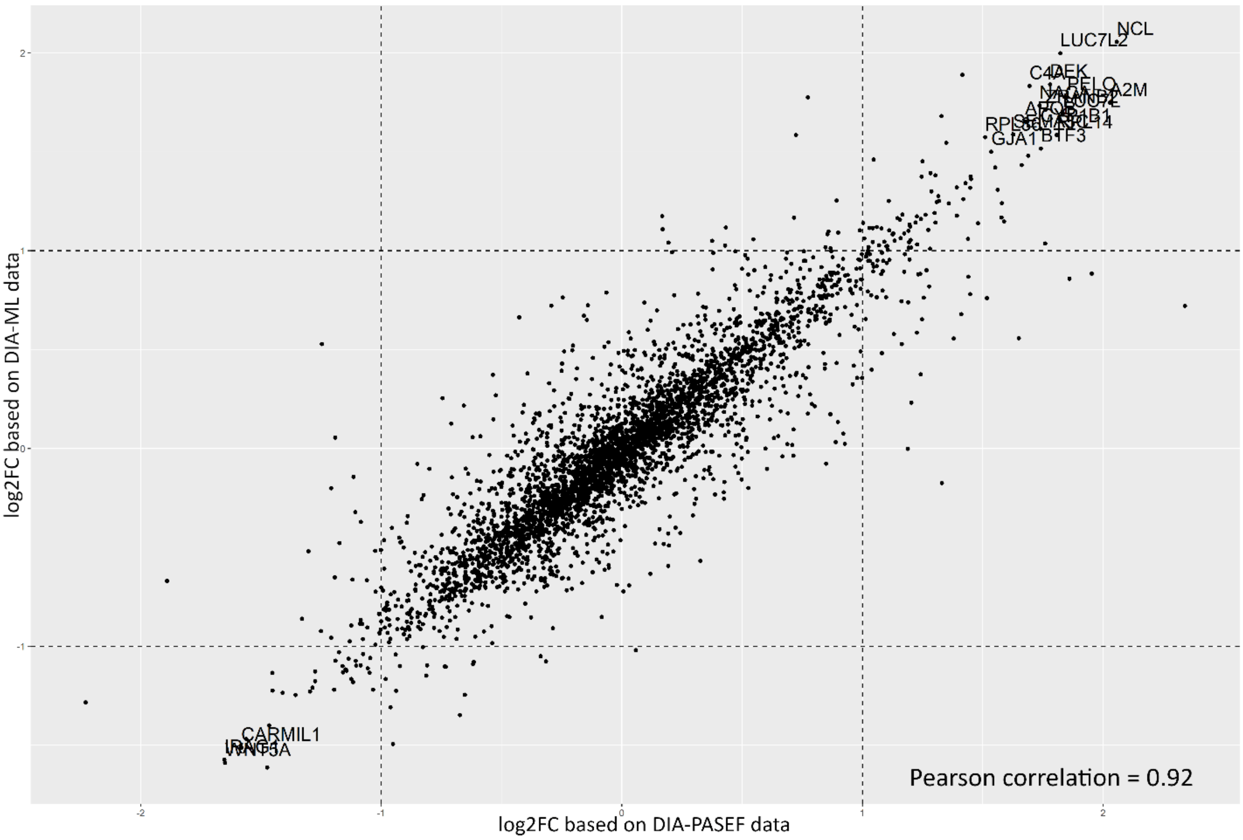


**Figure 12.** Correlation of Log2 Fold Changes for differentially expressed proteins in comparison of differentiated osteoblasts and differentiated VICs by DIA and DIA-ML proteomics.

**Supplementary Materials 3.** Clustering of osteoblasts and human valve interstitial cells (VICs) samples by principal component analysis (PCA).

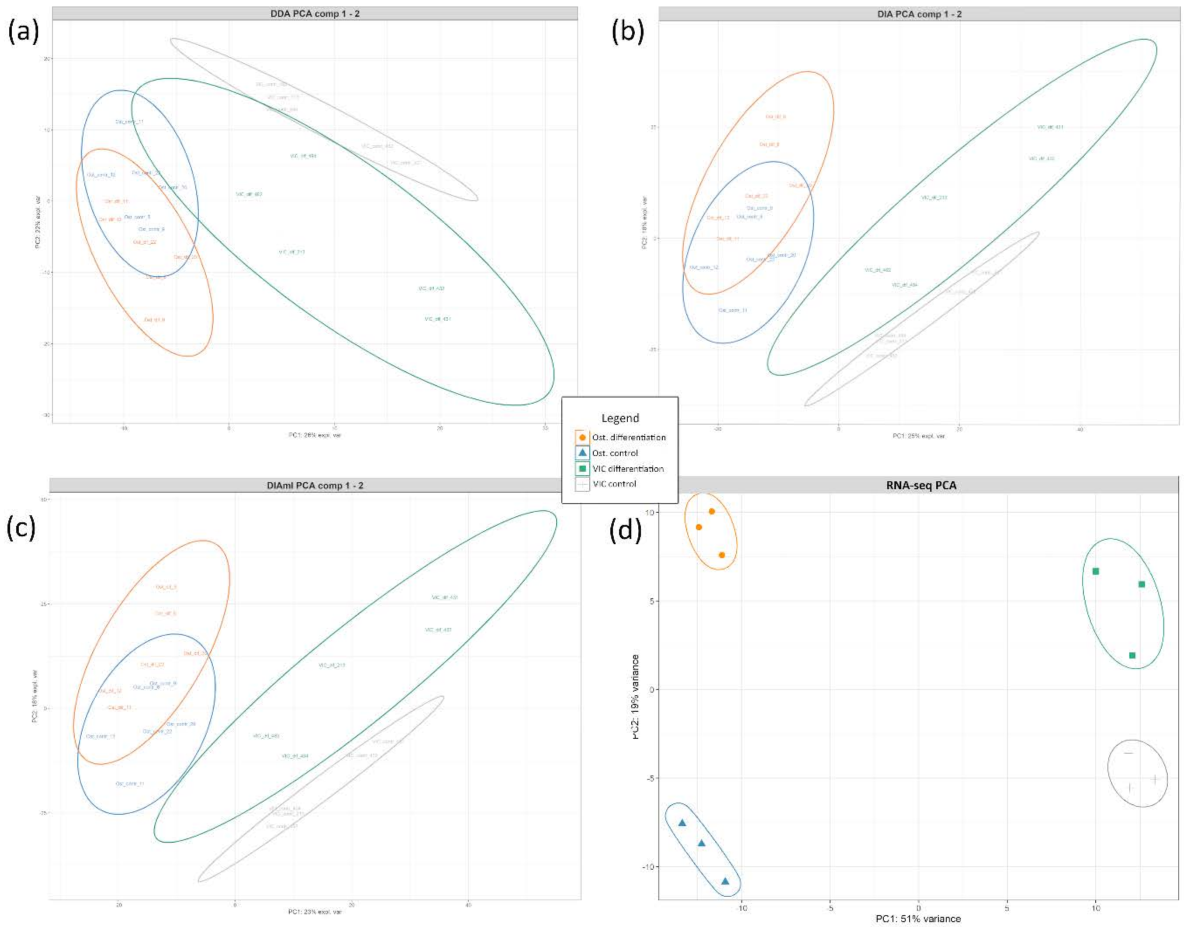


**Figure 13.** Clustering of human valve interstitial cells (VICs) and osteoblasts cultured in standard condition or after induction of osteogenic differentiation in the dimension of principal component analysis (PCA). (a) PCA clustering based on DDA-PASEF data. (b) PCA clustering based on DIA-PASEF data. (c) PCA clustering based on DIA-ML data. (d) PCA clustering based on RNA-seq data. Orange – osteoblasts on the 10th day after induction of osteogenic differentiation; Blue – osteoblasts in standard cultivation; Green – VICs on the 10th day after induction of osteogenic differentiation; Gray – VICs in standard cultivation.

**Supplementary Materials 4**. Venn diagrams representing the overlap of differentially expressed genes and proteins between control and osteogenic differentiation in osteoblasts and VICs (Valve Interstitial Cells).

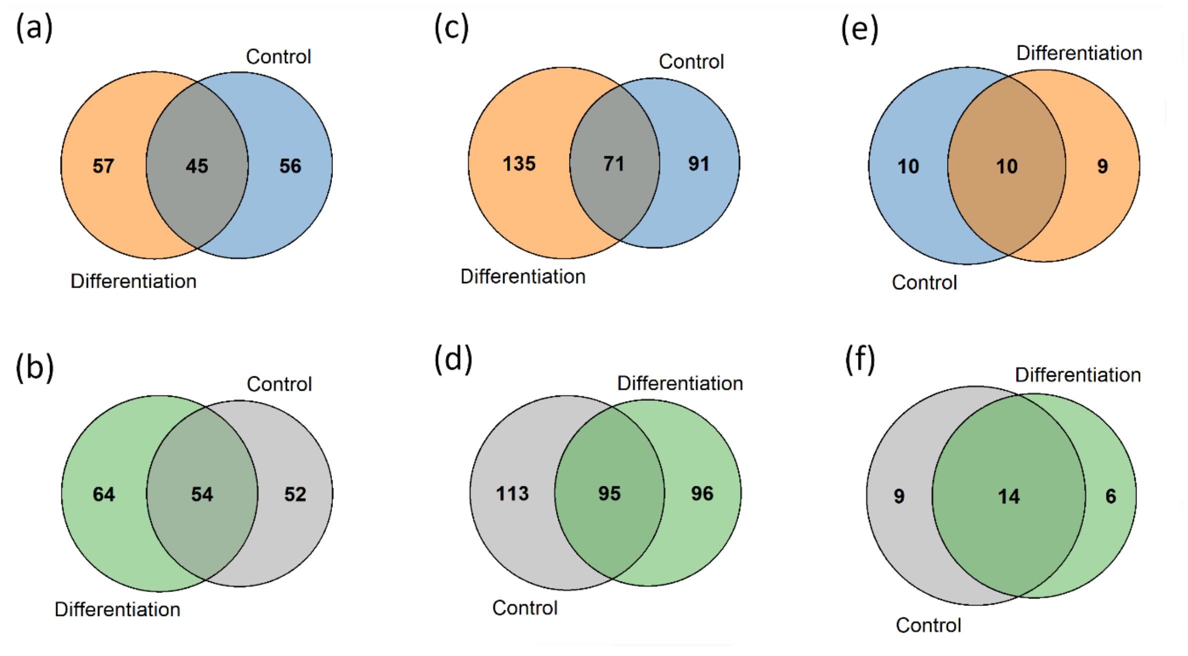


**Figure 14.** Venn diagrams representing the overlap of differentially expressed genes and proteins between control and osteogenic differentiation in (a, c, e) osteoblasts and (b, d, f) VICs. Overlaps are presented for (a, b) RNA-seq datasets, (c, d) Prot. w. datasets, and (e, f) Prot. rel. datasets. RNA-seq – transcripts identified by RNA-seq; Prot. w. – differentially expressed proteins, identified by at least one shotgun proteomics dataset; Prot. rel. – differentially expressed proteins, identified by all three shotgun proteomics datasets. Orange – osteoblasts on the 10th day after induction of osteogenic differentiation; Blue – osteoblasts in standard cultivation; Green – VICs on the 10th day after induction of osteogenic differentiation; Gray – VICs in standard cultivation.

**Supplementary Materials 5.** Boxplots for selective proteins are exclusively upregulated during VICs' osteogenic differentiation compared to osteoblasts differentiation.

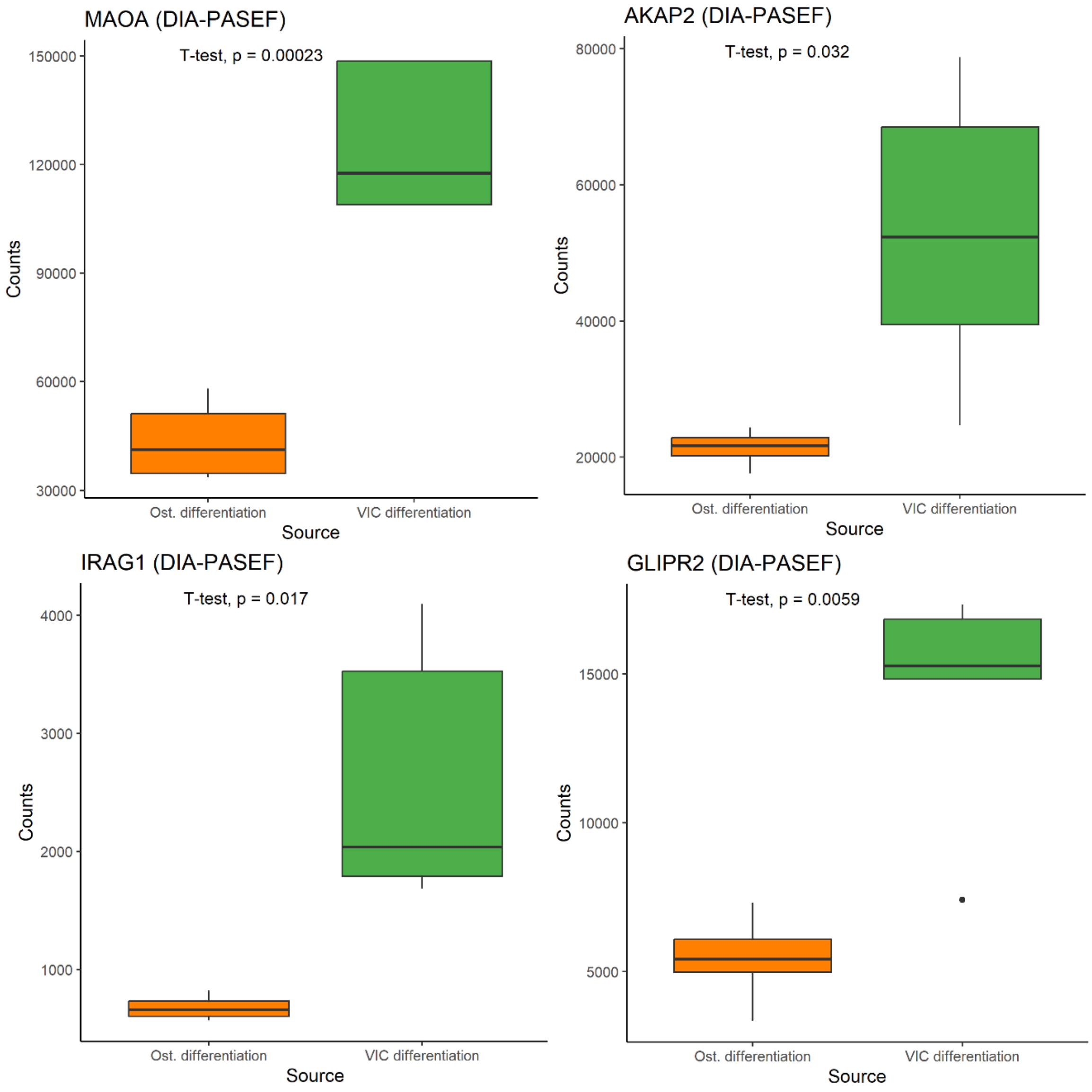


**Figure 15.** Proteins selectively upregulated in valve interstitial cell (VIC) differentiation compared to osteoblast differentiation, as indicated by the DIA-PASEF data. The x-axis represents the types of differentiated cells: orange - differentiated osteoblasts; green - differentiated VICs. The y-axis depicts the protein count numbers. The comparison was conducted utilizing a T-test, with statistical significance set at a p-value $< 0.05$.
